# An enzymatic–metabolic sensing axis in taste cells detects glucose-yielding carbohydrates

**DOI:** 10.64898/2026.03.13.710876

**Authors:** Aracely Simental-Ramos, Sandrine Chometton, A-hyun Jung, Amelia Cave, Thisun Udagedara, Caroline Metyas, Lilly Mai, Grace Tang, Jonathan Fan, Madeline Obrzut, Lindsey A. Schier

## Abstract

Glucose is a potent reinforcer of intake, yet most foods contain complex saccharides that do not yield free glucose until after digestion. How the oral sensory system rapidly evaluates the potential metabolic value of food remains unclear. Here, we identify an oral enzymatic–metabolic sensing mechanism that enables detection of glucose-yielding carbohydrates independent of canonical sweet taste receptors. Using genetic, virogenetic, molecular, and behavioral approaches in mice, we show that glucokinase (GCK) in taste cells is necessary for the attraction to glucose-containing sugars. We further demonstrate that maltase glucoamylase (MGAM), a glycosidic enzyme expressed on and near taste cells, facilitates rapid oral sugar sensing. Disruption of either GCK or MGAM in the major taste fields selectively attenuates the attraction to maltose and a carbohydrate-rich mixed diet, establishing both as intermediaries in the initial transduction pathway for complex saccharides that ultimately give rise to nutrient reward. Molecular profiling of taste papillae further revealed that deficient sweet sensing was accompanied by a compensatory increase in lingual MGAM, highlighting an adaptive mechanism for maintaining oral carbohydrate sensitivity. Together, these findings reveal that the oral epithelium actively preprocesses and metabolically evaluates dietary carbohydrates, providing a mechanism for rapid estimation of energetic value prior to ingestion.

**Significance Statement:** Carbohydrates strongly motivate eating, yet most are consumed in complex forms that do not immediately release glucose. We show that the oral epithelium contains an adaptive enzymatic–metabolic sensing system that enables rapid evaluation of glucose-yielding carbohydrates before digestion. Two enzymes, maltase glucoamylase and glucokinase, act locally in taste fields to preprocess and metabolically assess complex sugars, biasing ingestive behavior toward energetically favorable foods. This mechanism operates independently of canonical sweet taste receptors and is dynamically regulated by dietary experience and receptor sensitivity. These findings reveal that the mouth actively estimates the energetic value of food prior to ingestion, reshaping our understanding of how nutrient sensing guides dietary choice.

## Introduction

Carbohydrates are a primary source of metabolic energy and strongly motivate feeding behavior. Yet the energetic value of most dietary carbohydrates is not immediately accessible when food is first encountered. The majority of carbohydrates are consumed in complex forms that do not yield free glucose until after enzymatic digestion in the gut. This raises a fundamental question in nutrient sensing: how does the oral sensory system estimate the metabolic potential of food before it reaches the gut?

The taste system is traditionally viewed as the primary detector of the chemical quality of food, including sweetness. Mammalian taste buds contain multiple mechanisms for carbohydrate detection^1,2,3^. Monosaccharides (e.g., glucose, fructose) and disaccharides (e.g., sucrose, maltose) activate a G-protein–coupled receptor composed of T1R2 and T1R3 in type II taste cells; this receptor also responds to artificial sweeteners and certain D-amino acids^3,4,5,6^. Polysaccharides such as maltodextrins engage a distinct pathway whose receptor identity remains unresolved^2,7^. Signals from these receptors are transmitted to neural circuits involved in reward and homeostasis, ultimately promoting carbohydrate consumption^8^. However, the reliability of these sensory signals depends on their ability to predict the metabolic consequences of ingestion. When sweetness is dissociated from caloric value, as occurs with low-calorie sweeteners, neural and behavioral responses to sweet taste can weaken over time^9^.

Recent work suggests that glucose itself can be detected in the oral cavity through mechanisms independent of the canonical sweet receptor. Taste cells express specialized glucose-sensing machinery capable of responding directly to this key metabolic substrate^10^. Among these sensors is glucokinase (GCK), an enzyme that catalyzes the first step of glucose metabolism but also functions as a glucose sensor in several cell types, including pancreatic β-cells and neurons^12,13^. In the taste system, glucokinase is expressed in taste cells that lack the canonical sweet transduction machinery^10^. Activation of lingual GCK enhances neural responses in gustatory afferents and increases licking behavior toward glucose solutions^10^. Conversely, targeted loss of glucokinase in the major taste fields markedly reduces licking responses to glucose in both sweet-sensitive and sweet-blind mice^10^. These findings suggest that glucokinase-mediated signaling constitutes an oral metabolic sensor that contributes to the immediate valuation of glucose independently of sweet taste receptors.

Dietary experience can further shape the behavioral significance of this glucokinase-dependent sensory pathway. Repeated consumption of glucose and fructose produces a strong preference for the taste of glucose over fructose^14^. The more rewarding post-oral effects of glucose increase expression of Gck in taste buds, strengthening neural and behavioral responses to glucose^10^. Silencing Gck in the major taste fields, on the other hand, abolishes this learned sugar preference, demonstrating that dietary conditions can tune glucokinase-mediated sensory input to bias ingestion toward glucose^10^. Intriguingly, the same dietary experience also enhances attraction to maltose, a glucose–glucose disaccharide, relative to sucrose, which is composed of glucose and fructose^15^. This finding suggests a potential functional link between oral glucose sensing and the detection of complex carbohydrates that ultimately yield glucose. However, this hypothesis presents an important mechanistic challenge. In most foods, glucose is encountered as part of larger saccharides bound together by glycosidic linkages^11^. In this form, glucose cannot directly interact with intracellular metabolic sensors such as glucokinase. Therefore, if glucose-sensing mechanisms contribute to the oral detection of complex carbohydrates, an upstream process must first liberate free glucose within the taste environment.

Here we test the hypothesis that enzymatic processing of complex carbohydrates within the oral epithelium generates glucose that can engage glucokinase-dependent metabolic sensing pathways in taste cells (as well as the canonical sweet receptor pathways). Using genetic, virogenetic, molecular, and behavioral approaches in mice, we identify an enzymatic-metabolic sensing axis involving maltase glucoamylase (MGAM) and glucokinase (GCK) in the major taste fields. Our findings reveal that the activity of lingual enzymes involved in carbohydrate digestion and glucose metabolism enable sensitivity to dietary carbohydrates and bias ingestive behavior toward energetically favorable foods.

## Results

### Oral glucose detection relies on multiple pathways shaped by dietary experience

To date, the only known mechanisms for oral di- and poly-saccharide detection are found in a subset of taste cells (type II) that utilize the Transient Receptor Potential Cation Channel V (TRPM5)^16^. The taste of monosaccharides can also be signaled through this cellular pathway in type II cells, but glucose can engage other cell types and signaling pathways as well^17^. Glucokinase, for instance, is expressed in type III cells, which transduce taste information independently of TRPM5 (**Fig. 1a**). We thus first asked whether TRPM5-deficient mice are capable of discriminating glucose from isomolar fructose, and whether this ability depends on the dietary context. Transgenic Trpm5 mice were categorized as having sufficient (+) versus deficient (-) TRPM5-dependent taste signaling function (see Methods) and were compared to age-matched C57BL6/J (B6) mice (**Fig. 1b**). To our surprise, TRPM5- mice were also found to have a tendency towards lower expression of *Tas1r3*, a genetic marker for the sweet receptor, and lower expression of gustatory *Gck*, suggesting that constitutive Trpm5 may impact the expression of these respective upstream and parallel signaling intermediaries (**Fig. 1b**). Nevertheless, we compared the motivation to lick for glucose and fructose in a brief access taste test among sugar naïve (SN) TRPM5-, TRPM5+, and B6 mice (**Fig. 1c-d**). Consistent with our prior findings, naïve B6 mice licked in a similar concentration-dependent manner for both sugars (**Fig. 1d-e**); a comparable behavioral profile was observed for naïve TRPM5+ mice (**Fig. 1d-e**). In TRPM5- mice, on the other hand, licking for both sugars was relatively similar to water, and this response did not vary with concentration, suggesting impoverished monosaccharide sensitivity (**Fig. 1d-e**). As dietary experience with glucose and fructose vitalizes the attraction to glucose over fructose in both B6 mice and mice lacking the sweet receptor genes, we conditioned a separate cohort of B6, TRPM5+, and TRPM5- mice with daily access to glucose and fructose in the home cage for 24 days before the same brief access taste test (**Supplementary Fig. 1b**)^17^. Under these sugar exposure (SE) conditions, all groups displayed a preference for glucose, though the B6 and TRPM5+ mice clearly licked more avidly for glucose than the TRPM5- mice (**Fig. 1e-f**). The results show that although TRPM5 may both directly (via Type II cell signaling) and indirectly (via regulation of Tas1r3 and *Gck*) contribute to oral glucose sensing, residual signaling via a TRPM5-independent mechanism confers some sensitivity, and this can be enhanced by dietary conditions.

**Fig. 1:**
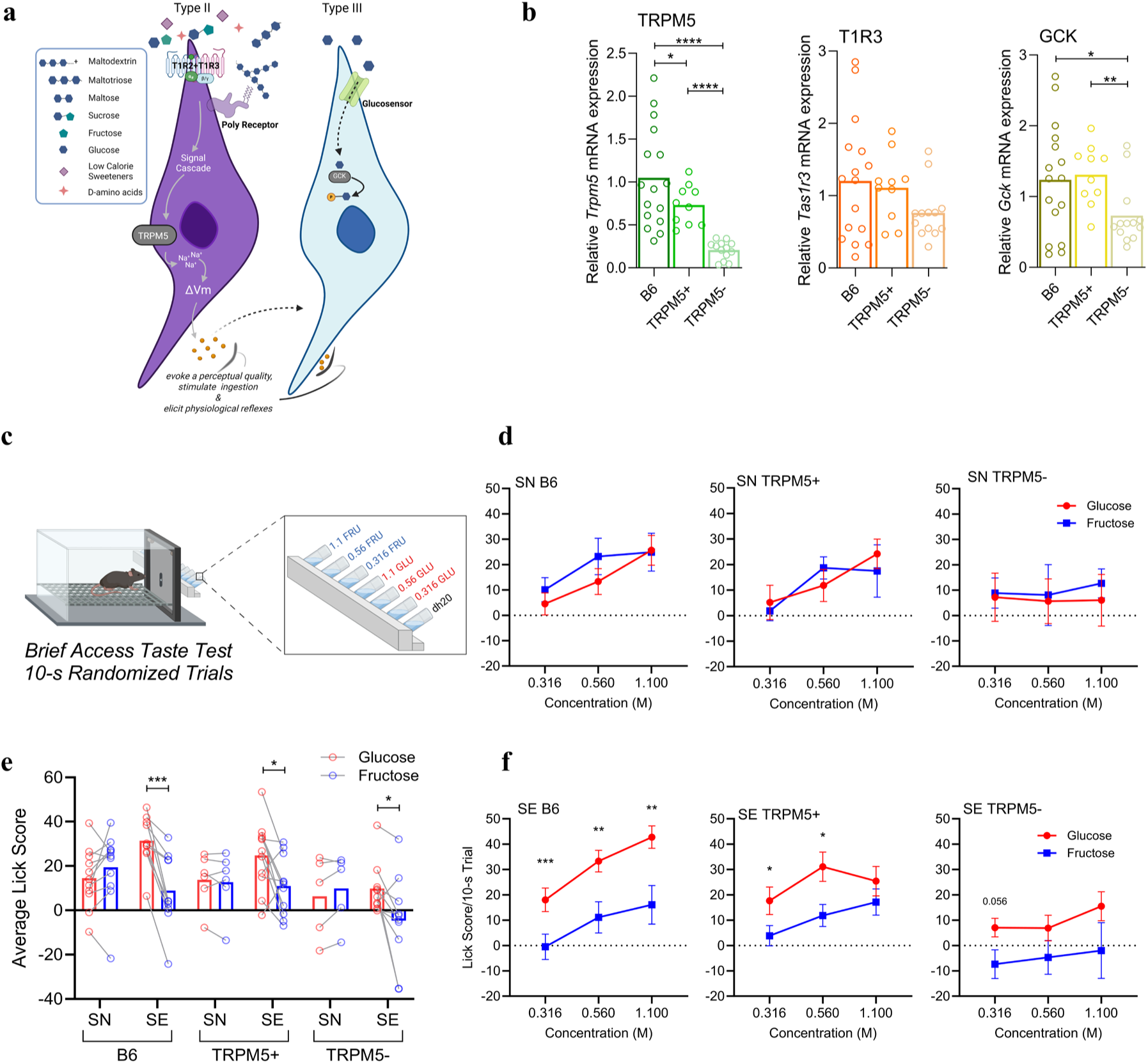
TRPM5 deficient and sweet sensitive mice lick more avidly for glucose than for fructose with sugar exposure. **a** Hypothetical schematic of Type II and Type III sweet taste cell. **b** Mean Trpm5, Tas1r3, and Gck relative transcript expression in the circumvallate taste tissue of sugar exposed and sugar naive B6 (n =16), TRPM5+ (n=10-11), and TRPM5- (n=13). **c** Schematic depicting glucose vs fructose brief access taste test in lickometer. **d** Mean (±SEM) lick scores of 0.316M, 0.56M and 1.1M glucose and fructose in brief access taste test for sugar naïve (SN) B6 (n=11), TRPM5+ (n=6) and TRPM5- (n=5). **e** Mean average lick score for B6 (n=21), TRPM5+ (n=17) and TRPM5- (n=15). **f** Mean (±SEM) lick score of 0.316M, 0.56M and 1.1M glucose and fructose in brief access taste test for sugar exposed (SE) B6 (n=10), TRPM5+ (n=11) and TRPM5- (n=10). (*:p<0.05, **:p<0.01, ***:p<0.001,****:p<0.0001). All tests were conducted in the gustometer. Statistical Analysis are in Supplementary Table 1.

### Glucokinase drives maltose attraction independent of canonical taste-signaling pathways

We next assessed how the naïve and sugar-exposed mice respond to two glucose containing disaccharides, maltose and sucrose (**Fig. 2a**). As expected, naïve B6 and TRPM5+ licked more for sucrose (**Fig. 2b-c**). This pattern is consistent with the fact that sucrose is more effective at engaging the sweet receptor than maltose^18^. Naïve TRPM5- mice, on the other hand, were equally responsive to both sugars, in a concentration-dependent manner and well above the water baseline (**Fig. 2b-c**). Dietary conditioning with glucose and fructose revised these profiles in all three groups. Specifically, both B6 and TRPM5+ mice increased their licks for maltose, to levels that were more comparable to the sucrose (**Fig. 2c-d**), and TRPM5- mice continued to lick for maltose while selectively reducing licking for sucrose (**Fig. 2c-d**). These results confirm our earlier findings that oral sensitivity to glucose can generalize to maltose, at least under certain dietary conditions, and this does not depend on the canonical carbohydrate sensing pathway involving TRPM5^15^.

**Fig. 2:**
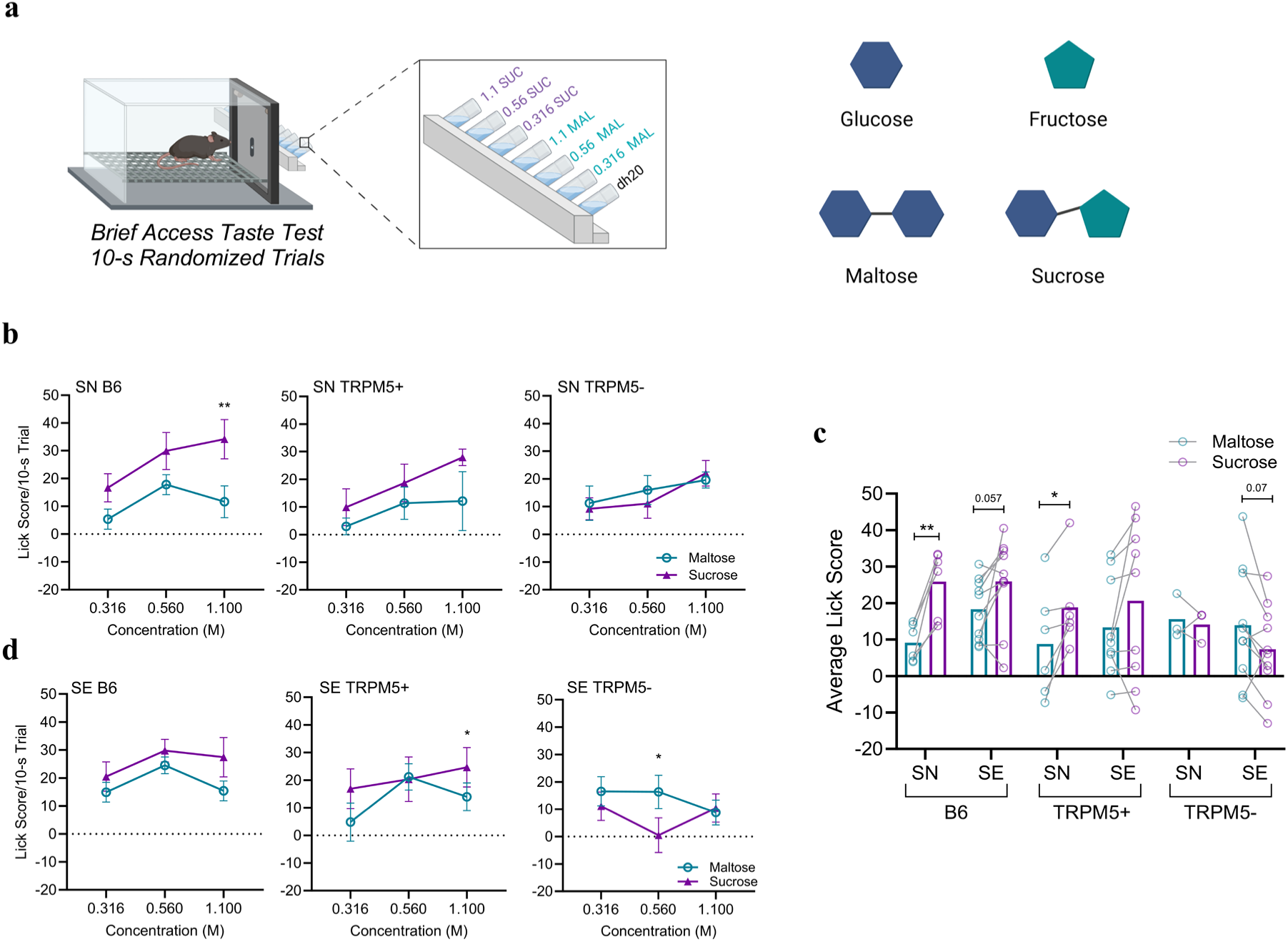
TRPM5 deficient and sweet sensitive mice lick more avidly for maltose than sucrose after sugar exposure and show differing levels of taste genes mRNA expression. **a** Schematic depicting maltose vs sucrose brief access taste test in lickometer and schematic showing representative depictions of glucose, fructose, maltose and sucrose. **b** Mean (±SEM) lick score of 0.316M, 0.56M, and 1.1M maltose and sucrose in brief access taste test for sugar naive (SN) B6 (n=8), TRPM5+ (n=6) and TRPM5- (n=3). **c** Mean average lick score across concentrations for maltose and sucrose in B6 (n=16), TRPM5+ (n=15), TRPM5- (n=13). **d** Mean (± SEM) lick score of 0.316M, 0.56M, and 1.1M maltose and sucrose in brief access taste test for sugar exposed (SE) B6 (n=10), TRPM5+ (n=9) and TRPM5- (n=10). (*:p<0.05, **:p<0.01). All tests were conducted in the gustometer. Statistical Analysis are in Supplementary Table 2.

In sweet-sensitive mice, habitual intake of glucose and fructose drives expression of *Gck* in the taste buds and ultimately bestows a heightened attraction to glucose^10^. After confirming that Trpm5 deficient mice also display this heightened attraction to glucose containing sugars, we next asked whether GCK contributes to the appeal of maltose. A separate cohort of TRPM5+ and TRPM5- mice were provided daily exposure to glucose and fructose for 18 days (**Supplementary Fig. 1c**). After this, we used a shRNA to acutely silence *Gck* in the major taste fields or delivered a scrambled control treatment (**Fig. 3a**). This approach reduces *Gck* in the taste papillae by ∼ 40%^10^. Five days later mice were tested for their relative preference for maltose and sucrose in a brief access taste test (**Fig. 3b**). In TRPM5+ mice, loss of lingual *Gck* led to a significant reduction in the motivation to lick for maltose, without impacting the motivation for sucrose (**Fig. 3c-d**). In TRPM5- mice, loss of lingual GCK also reduced the motivation to lick for maltose (**Fig. 3e-f**). The results provide the first evidence that *Gck*-mediated signaling drives the attraction to maltose, and this can occur independently of canonical type II cell TRPM5-dependent input.

**Fig. 3:**
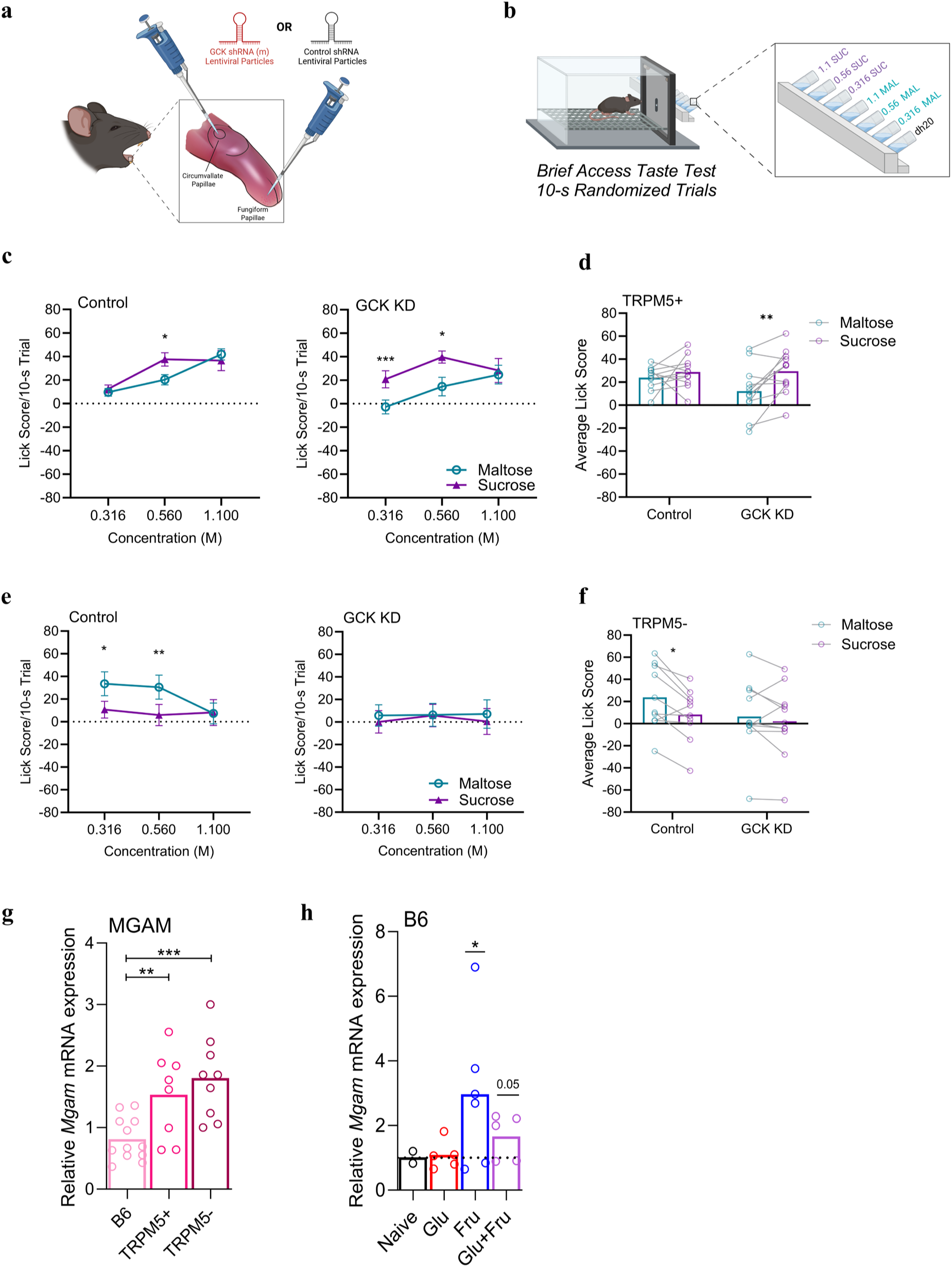
Lingual glucokinase virogenetic silencing impairs behavioral sensitivity to maltose. **a** Schematic of Gck virogenetic shRNA or control shRNA silencing surgery in the major taste fields of the tongue. **b** Schematic of brief access taste test of maltose and sucrose in lickometer. **c** Mean (±SEM) lick score of 0.316M, 0.56M, 1.1M maltose and sucrose in control (n=11) and GCK KD (n=12) TRPM5+. **d** Mean lick scores averaged across concentration for control and GCK KD TRPM5+ (11-12/group). **e** Mean (±SEM) 0.316M, 0.56M, 1.1M maltose and sucrose in control (n=10) and GCK KD (n=11) TRPM5-. **f** Mean lick scores averaged across concentration for control and GCK KD TRPM5- (10-11/group). **g** Mean relative transcript expression of Mgam for B6 (n=8), TRPM5+ (n=8) and TRPM5- (n=9). **h** Mean (±SEM) relative MGAM transcript for B6 sugar naïve (n=2), glucose experience (n=5), fructose experience (n=6), or glucose + fructose experience(n=5). (*:p<0.05, **:p<0.01,***:p<0.001,****:p<0.0001). All tests were conducted in the Davis Rig. Statistical Analysis are in Supplementary Table 3.

### Enzymatic preprocessing of maltose at the taste cells enables GCK-mediated sensing of this complex sugar

These compelling findings present a quandary though, as the glucose molecules in maltose are bound together by an α(1→4) linkage and, therefore, cannot directly interact with GCK in this form^19^. This implies that there is a complementary system upstream of the GCK-linked sensor that can rapidly generate free glucose from the disaccharide. Amylase, a digestive enzyme that can break down polysaccharides into di- and tri-saccharides is present in the saliva^20^. However, amylase cannot further digest maltose into free glucose. However, published studies have recently identified α-glucosidases, such as maltase glucoamylase (MGAM) and sucrase isomaltase, in the apical membranes of taste cells^21^. Broad spectrum pharmacological inhibition of these enzymes attenuates responsivity of the primary taste nerve to disaccharides but whether this is sufficient to impact behavioral sensitivity remains unclear^21^. As MGAM is responsible for the majority of maltose digestion in the gut, we hypothesized that lingual MGAM may also play an important role in the maltose taste preference^1^.

To investigate this possibility, we quantified *Mgam* expression in B6, TRPM5+ and TRPM5- mice and observed that MGAM was higher in the TRPM5+ and TRPM5-mice compared to B6 **(Fig. 3g**). To see if Mgam levels are regulated by dietary sugar, we compared relative Mgam expression levels in B6 mice that were kept naïve, provided regular access to just glucose, just fructose, or both glucose and fructose. These analyses revealed that mice with a history of consuming both glucose and fructose or just fructose especially displayed significantly greater *Mgam* expression in the taste papillae, while a history of consuming just glucose did not drive *Mgam* expression (**Fig. 3h**). The presence of free glucose does not necessitate the upregulation of *Mgam*, while experience with other sugars leads to its higher expression. The dynamic presence of Mgam in taste cells corroborates the theory of its contribution to carbohydrate digestion.

Further, since chronic exposure to glucose and fructose can enhance the relative attraction to maltose (vs sucrose), we asked whether dietary exposure to the two disaccharides could likewise sufficiently condition a heightened attraction maltose, and favorably re-program *Gck* and *Mgam* expression in the taste buds of sweet-sensitive mice (**Supplementary Fig.1e-f**). Indeed, B6 mice exposed to these sugars displayed a heightened avidity for maltose, combined with a reduced avidity for sucrose during a brief access taste test (**Fig. 4a-c**). Interestingly, this avidity to maltose did not transfer to a glucose preference (**Fig. 4d-f**). However, the behavioral changes were associated with an upregulation of *Gck* and *Mgam* with no effect on *Tas1r3* levels (**Fig. 4g**). Based on these collective results, we reasoned that greater *Mgam* levels may facilitate sensing of sugar when other key signaling intermediaries are impoverished [(i.e., TRPM5 in the TRPM5+ and – mice, see **Fig. 3g**); *Tas1r3* and *Gck* in the TRPM5- mice, see **Fig. 1b**)], when glucose resources are scarce in the environment (fructose only condition, **Fig. 3h**), or when nutritional conditioning reprograms oral sensitivity towards dietary sugars that ultimately yield more glucose (**Fig. 4a-c**). In other words, we posit that enzymatic activity in the major taste fields generates more ligands (as free glucose) for nearby receptors and empowers rapid detection of favorable sources of energy (see hypothetical model in **Fig. 4h**), with Mgam itself regulated by both genetic and environmental factors.

**Fig. 4:**
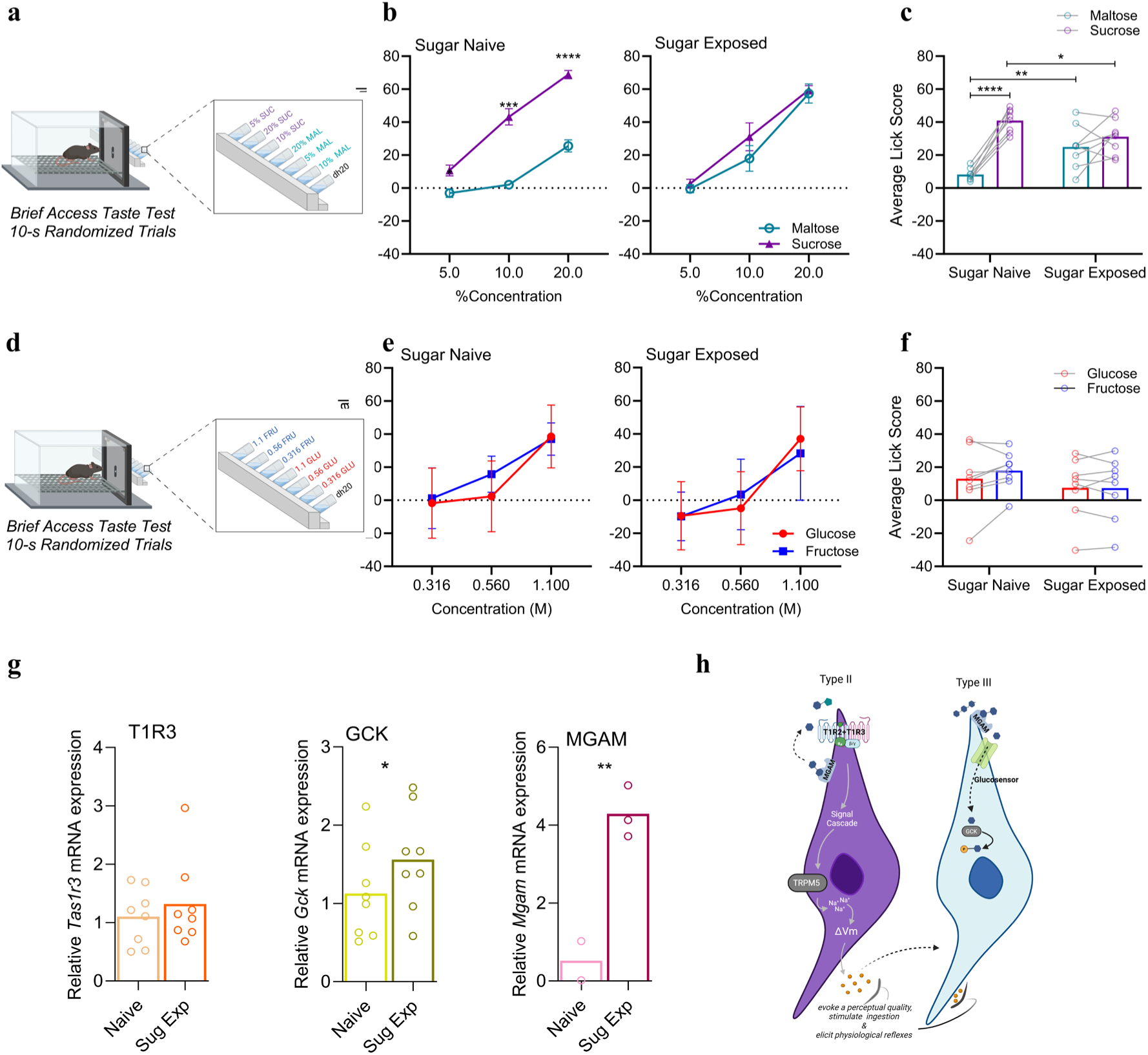
Prolonged exposure to maltose and sucrose elevates licking for maltose and glucosensing enzyme expression. **a** Schematic depicting maltose vs sucrose brief access taste test in lickometer. **b** Mean (±SEM) lick score for 5.0%, 10.0% and 20.0% maltose and sucrose for sugar exposed and sugar naive (n=8/group). **c** Mean lick score for maltose and sucrose averaged across concentration (n=8/group). **d** Schematic depicting glucose vs fructose brief access taste test in lickometer. **e** Mean (±SEM) lick score for 0.316M, 0.56M, and 1.1M glucose and fructose in brief access taste test for sugar naive B6 (n=8) and sugar exposed B6 (n=8). **f** Mean average lick score across concentrations for sugar naive and sugar exposed B6 (n=8/group). **g** Mean relative transcript expression of Tas1r3, Gck, and Mgam in sugar naive (n=2-8) and sugar exposed (n= 3-8). (*:p<0.05, **:p<0.01, ***:p<0.001,****:p<0.0001). **h** Schematic of Type II and Type III sweet taste cells depicting MGAM and GCK function. All tests were conducted in the Davis Rig. Statistical Analysis are in Supplementary Table 4

### Sweet-subsensitive mice exhibit higher levels of *Mgam* in the taste papilla and lick avidly for maltose

If our hypothesis is correct, then diminished sweet receptor function should also be compensated for by greater digestive capacity in the taste cells. Thus, we assessed circumvallate *Mgam* levels in mice (of the 129 strain) that carry a variant in the *Tas1r3* gene which renders the receptor less effective at binding sweeteners compared to mice that carry the sweet-sensitive Tas1r3 allele (B6) (**Fig. 5a**)^22,27^. Strikingly, naïve 129 mice had significantly greater *Mgam* in the taste papilla than B6 mice (**Fig. 5b**). The 129 had significantly less Tas1r3, but similar amounts of *Gck*, versus the B6 mice (**Fig. 5b**). To test if this 129 profile permitted sufficient sensitivity to maltose, we measured licking responses to a wide range of maltose concentrations in naïve mice of each strain (**Fig. 5c**). The data showed that 129 mice licked in a normal concentration-dependent manner for maltose (**Fig. 5d**). These results parallel our findings in mice that are rendered subsensitive through a different mechanism (loss of TRPM5) and collectively suggest that the ability to rapidly cleave complex sugars to free glucose may compensate for reduced receptor function.

**Fig. 5:**
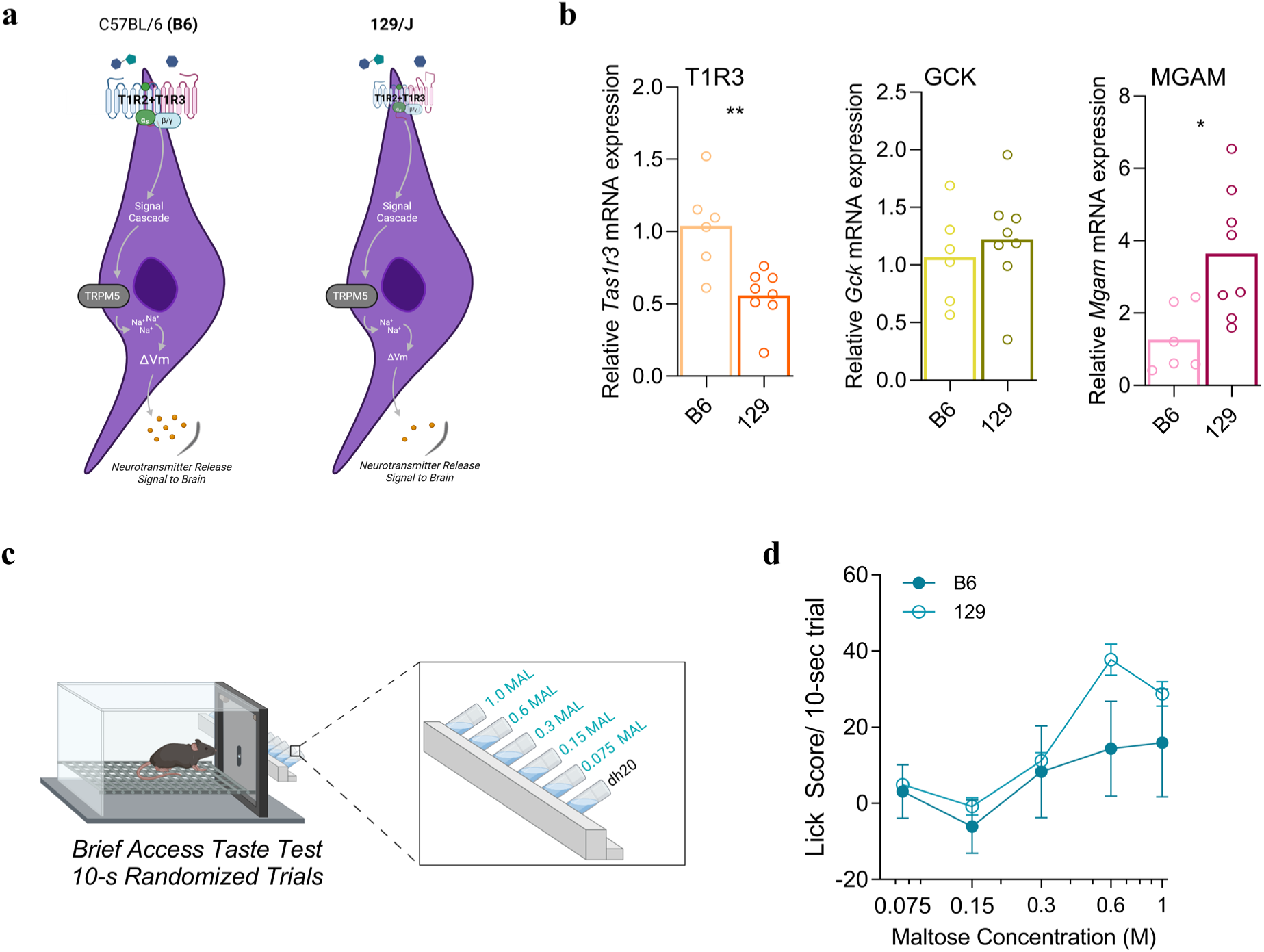
Sweet subsensitive mice (129/J) lick as avidly for maltose as sweet sensitive mice and have elevated expression of Maltase-Glucoamylase. **a** Schematic depicting B6 and 129/J representative taste cells showing Tas1r3 polymorphism in 129/J. **b** Mean relative transcript expression of Tas1r3, Gck and Mgam in the circumvallate taste tissue of B6 (n= 6) and 129 (n=6) groups. **c** Schematic of brief access taste test for 0.075M, 0.15M, 0.3M, 0,.6M and 1.0M maltose concentrations and water. **d** Mean (±SEM) maltose brief access taste test lick scores for B6 (n=3) and 129 (n=3). (*:p<0.05, **:p<0.01). All tests were conducted in the gustometer. Statistical Analysis are in Supplementary Table 5.

### Loss of *Mgam* in the major taste fields reduces the hedonic value of maltose

No study has directly tested if MGAM within the taste fields plays a meaningful role in sugar’s appeal. To test this, we developed a virogenetic approach to selectively silence *Mgam* in the major taste fields using an shRNA knockdown virus in TRPM5+ mice (**Fig. 6a**). This resulted in a ∼40-50% knockdown of *Mgam*, without affecting *Gck* or *Tas1r3* expression (**Fig. 6b**). We utilized this method in a different cohort of mice, then five days after shRNA (or scrambled control) treatment, we subjected naïve TRPM5+ mice to a short licking test for 0.6 M maltose and analyzed the microstructural patterns of licking (**Fig. 6c-f**). Prior work has shown that the number of licks in a burst of licking, which is defined as a period of continuous licks each separated by less than 1 second, increases with the hedonic value of the tastant (**Fig. 6e**)^23^. We found that knockdown of *Mgam* in the major taste fields resulted in significantly smaller bursts and thus required the mice to take more bursts to reach the 300-lick criterion, as compared to the control treatment (**Fig. 6f**). These results provide the first evidence that peri-taste MGAM potentiates the rapid detection and ingestion of sugar.

**Fig. 6:**
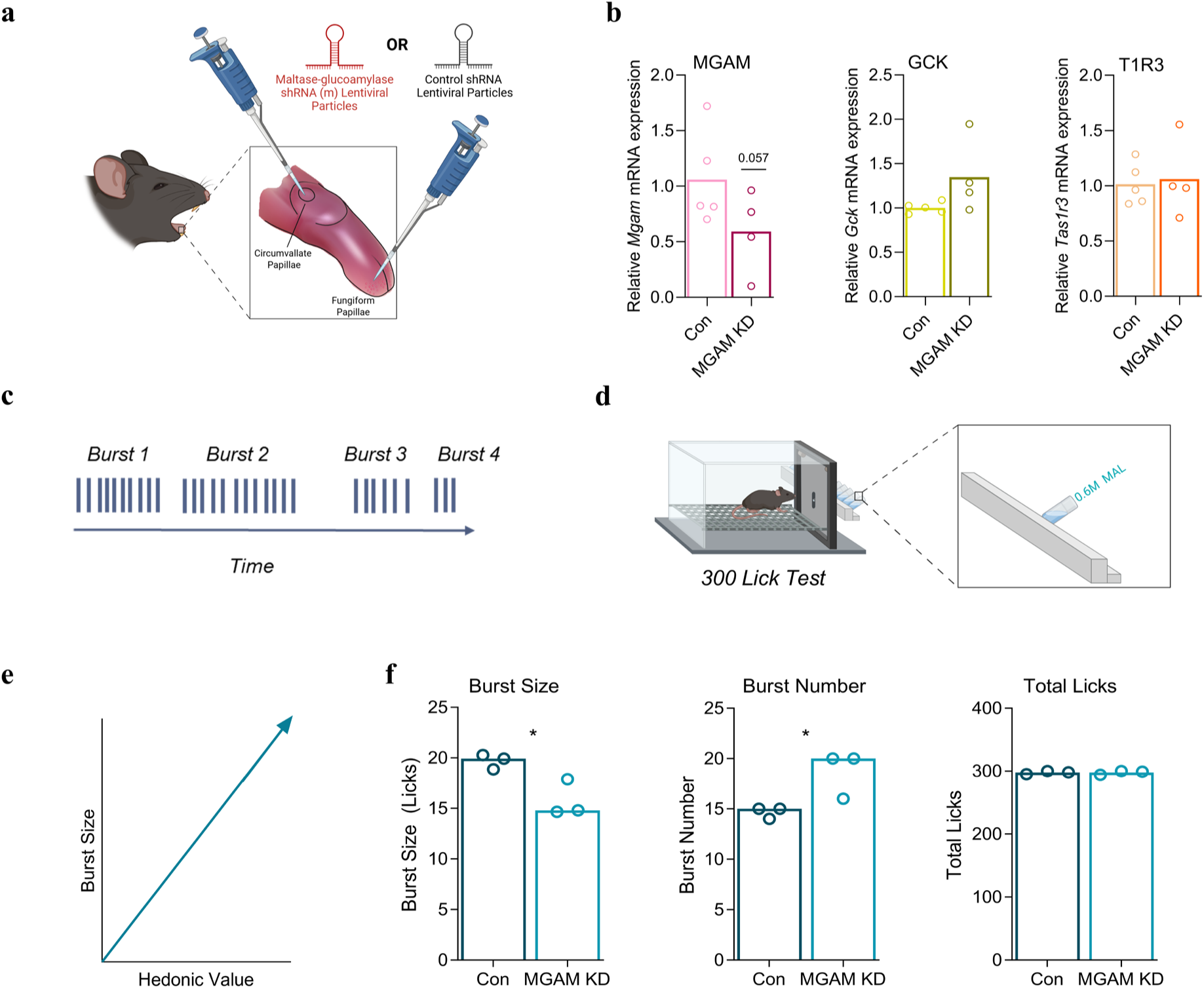
Lingual maltase-glucoamylase virogenetic silencing reduces hedonic value of maltose. **a** Schematic of control and Mgam virogenetic shRNA silencing surgery in the major taste fields of the tongue. **b** Mean relative transcript expression of Mgam, Gck, and Tas1r3 in control (n= 4) and MGAM KD (n=4) in TRPM5+ mice **c** Schematic of burst structure (period of continuous licks separated by <1s) and time frame **d** Schematic of 300 lick test of 0.6M maltose in lickometer. **e** Schematical graph showing relationship between burst size and hedonic value. **f** Mean (±SEM) burst size, burst number, total licks for 300 lick test in control (n=3) and MGAM KD (n=3) in TRPM5+ mice (*:p<0.05). All tests were conducted in the gustometer. Statistical Analysis are in Supplementary Table 6.

### Loss of *Mgam* in the major taste fields reduces the affinity for maltose in sugar-exposed mice

We also sought to determine whether MGAM contributes to the acquired appeal of maltose through regular sugar exposure independently of sweet taste using mice lacking the *Tas1r2* and *Tas1r3* genes (T1R KO). In our recently published study, we found that naïve T1R KO mice had more *Gck* in the taste papillae compared to naive B6 mice, while sugar-exposed T1R KO had less *Gck* in the taste papillae than sugar-exposed B6 mice; we did not directly compare across exposure conditions within each strain ^48^. Thus, here we first confirmed that T1R KO with a history of prolonged access to glucose and fructose in the home cage, displayed upregulated *Mgam* expression, and, to a lesser extent, *Gck* expression. These results further demonstrate that sweet receptor signaling is not required for diet-driven changes in the expression of these enzymes in the taste papillae (**Fig. 7a-b**). Replicating our previous studies, in a sperate cohort of T1R KO mice, we also found that sugar-exposed T1R KO mice lick substantially more for glucose relative to fructose by the end of the sugar exposure paradigm (**Supplementary Fig. 1d)**. We then performed the virogenetic knockdown of *Mgam* (or its scrambled control) and measured licking responses for maltose and sucrose five days later (**Fig. 7c-d**). Overall, we found that sugar-exposed T1R KO mice did not lick very avidly for these two sugars; they did, however, lick more for maltose than sucrose at the lowest concentration tested (**Supplementary Fig. 3a**). This preference was eliminated when *Mgam* was selectively knocked down in the major taste fields (**Supplementary Fig. 3a-b**). Upon close inspection of the data, we noticed that licks for water were relatively high on this test (control: 24.32 ± 7.05; MGAM KD: 26.80 ± 4.36). This led us to wonder if the high concentrations of sugar presented in this test saturated the enzymatic and sensing mechanisms of these sweet-blind mice as trials progressed; such an effect would interfere with the saliency of each sugar within the trial it was presented. Thus, we separately analyzed licking responses to the very first trial with each sugar on this test (referred to as ‘Block 1’). The results revealed a more robust preference for maltose (versus sucrose) in the control group, which was again largely driven by a significant elevation in licking for maltose at the lowest concentration tested (Block 1 water licks for control: 8.0 ± 4.59) (**Fig. 7e-f**). This effect was notably absent from the MGAM KD group. Instead, this group licked comparably for sucrose and maltose at all concentrations tested (water licks for MGAM KD: 16.80 ± 1.36). To confirm that these effects were specific to the disaccharide, we tested all mice with glucose and fructose (for which MGAM is not needed). As expected, MGAM KD did not interfere with glucose preference, for neither the first block nor all the combined blocks, confirming the specificity of this manipulation (**Fig. 7g-i; Supplementary Fig.3c-d)**.

**Fig. 7:**
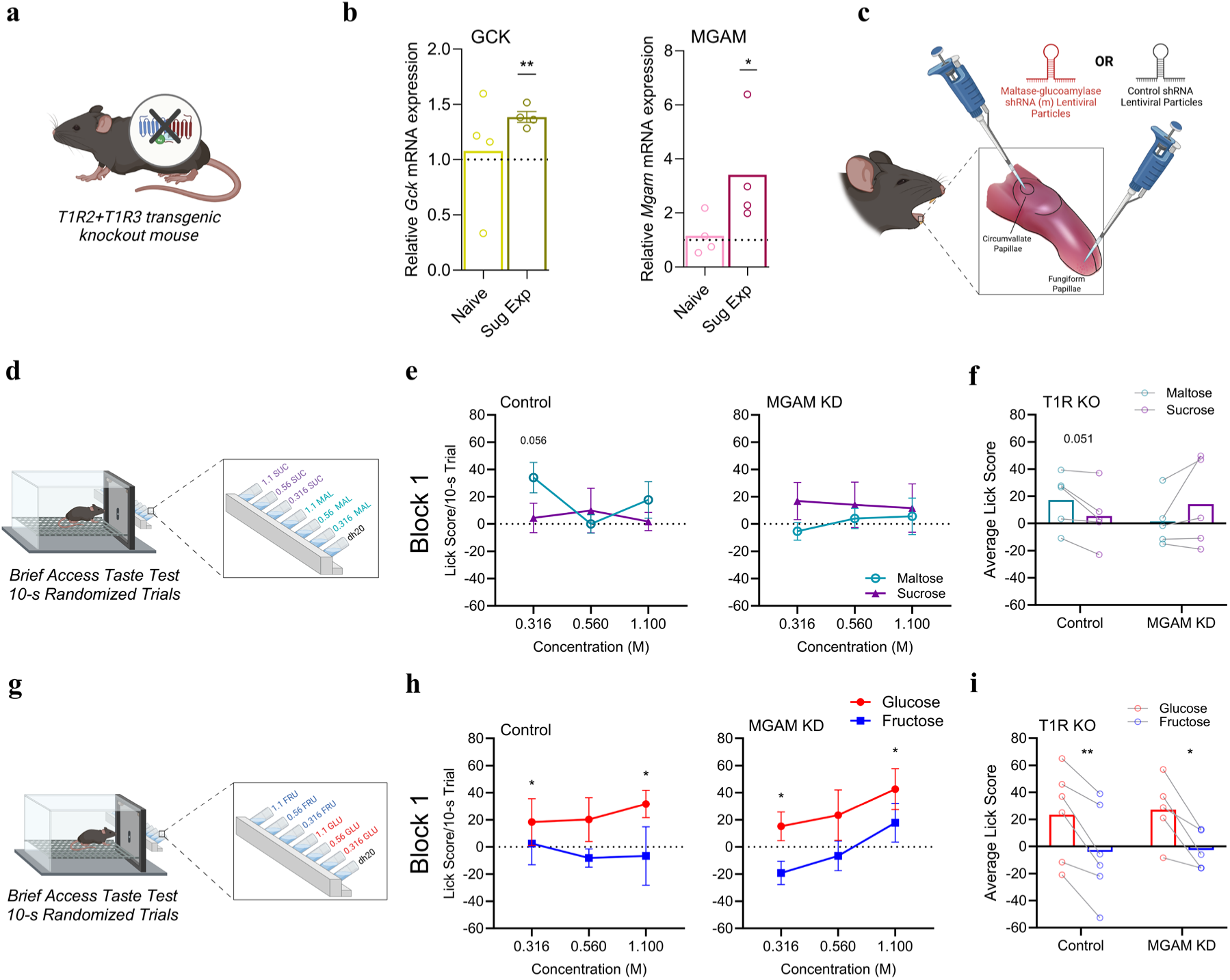
Lingual silencing of maltase-glucoamylase impairs sensitivity to maltose but not glucose in sweet receptor knockout mice. **a** Schematic depicting T1R2+T1R3 sweet receptor transgenic knockout mouse. **b** Mean transcript expression of Gck and Mgam for sugar naive (n=4) and sugar exposed (n=4) T1RKO. **c** Schematic of control and Mgam virogenetic shRNA silencing surgery in the major taste fields of the tongue. **d** Schematic of brief access taste test of maltose and sucrose in lickometer. **e** Block 1 mean (±SEM) lick scores for 0.316M, 0.56M, 1.1M maltose and sucrose in control (n=5) and MGAM KD (n=5) T1RKO. **f** Mean lick scores averaged across concentration for control and MGAM KD for first block (5/group). **g** Schematic of brief access taste test of glucose and fructose in lickometer. **h** First block mean (±SEM) lick scores for 0.316M, 0.56M, 1.1M glucose and fructose in control (n=6) and MGAM KD (n=5) T1RKO. **i** First block glucose and fructose mean lick scores averaged across concentration for control (n=6) and MGAM KD (n=5) T1RKO. All tests were conducted in the Davis Rig. (*:p<0.05, **:p<0.01, ***:p<0.001, ****:p<0.0001). Statistical Analysis are in Supplementary Table 7.

### Silencing of *Mgam* or *Gck* reduces initial avidity to a carbohydrate-rich diet

Processed foods, including ultra processed foods, comprise more than half of the average Western diet^24^. These foods are high in simple sugar and refined starches like maltodextrins. Thus, in a final experiment, we asked whether GCK and MGAM contribute to the innate tastiness of a mixed nutrient food. Peptamen is a liquid diet that is 33% fat, 12% protein, and 55% carbohydrate, with all of the carbohydrate content coming from maltodextrins and starches (contains no mono- or di-saccharides)^25^. First, we characterized the microstructural licking patterns for Peptamen in a 1000 lick test in naïve B6 mice (**Fig. 8b**, pre-test). Most of the mice easily reached the lick criterion within 20 minutes (**Fig. 8c-d**). Next, the mice were split into three groups and received either the *Gck* shRNA (GCK KD), *Mgam* shRNA (MGAM KD), or scrambled sequence shRNA (control) treatment to the major taste fields (**Fig. 8a**). Five days later, we measured licking patterns to the same Peptamen solution in a 1000 lick post-test (**Fig. 8b**). Generally speaking, all groups licked more avidly for the diet on this second exposure, with mice licking in larger bursts, indicating an increased hedonic appeal, and reaching the 1000 lick criterion more rapidly on the post-test (**Figures 8d-f**). Since both oral and post-ingestive factors may contribute to this acquired appetition within the 1000 lick test, we additionally analyzed how licking progressed across the early part of the meal, the time during which intake behaviors are primarily controlled by orosensory factors We ascertained that whereas the mixed diet stimulated a robust appetition response in the control group during the post-test, it failed to propel this incipient response in mice with diminished *Gck* or *Mgam* in the major taste fields (**Fig. 8g**). These findings reveal that non-canonical oral processes associated with carbohydrate digestion and metabolic signaling are necessary to swiftly engage ingestive behaviors that promote intake of complex carbohydrates in a mixed nutrient medium.

**Fig. 8:**
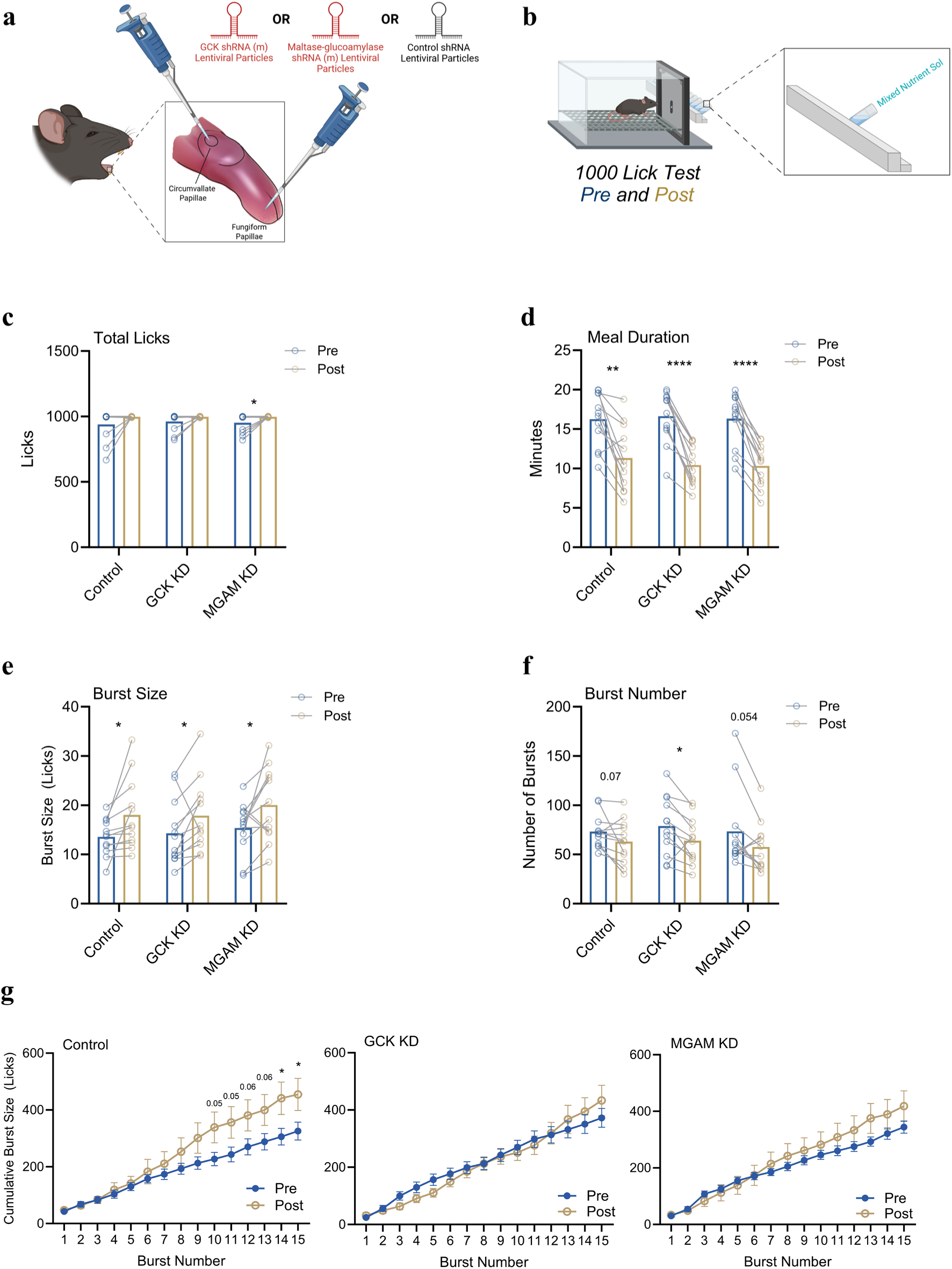
Lingual virogenetic silencing of glucokinase or maltase-glucoamylase impairs initial detection of mixed nutrient solution (Peptamen). **a** Schematic showing control, Gck, and Mgam virogenetic shRNA silencing surgery in the major taste fields of the tongue. **b** Schematic of 1000 lick test of mixed nutrient solution (Peptamen) in lickometer for pre-surgery and post-surgery. **c** Mean total licks for 1000 lick test in control, GCK KD, MGAM KD pre-test vs post-test. **d** Mean meal duration for 1000 lick test in control, GCK KD, MGAM KD pre-test vs post-test. **e** Mean burst size for 1000 lick test in control, GCK KD, MGAM KD pre-test vs post-test. **f** Mean burst number for 1000 lick test in control, GCK KD, MGAM KD pre-test vs post-test. **g** Mean (±SEM) cumulative first 15 burst size for 1000 lick test in control, GCK KD, MGAM KD pre-test vs post-test. (n=12/group for all tests) (*:p<0.05, **:p<0.01, ***:p<0.001, ****:p<0.0001). All tests were conducted in the gustometer. Statistical Analysis are in Supplementary Table 8.

Together, the findings identify a previously unrecognized enzymatic-metabolic sensing axis within the oral epithelium. Maltase glucoamylase generates free glucose from complex saccharides in taste fields, enabling activation of glucokinase-dependent metabolic sensors that bias ingestion toward glucose-yielding carbohydrates.

## Discussion

Efficient feeding requires organisms to estimate the energetic value of food before ingestion, despite encountering nutrients in chemically diverse and often metabolically inaccessible forms. Although carbohydrates robustly engage sensory–reward circuits, the mechanisms by which the oral sensory system predicts the metabolic consequences of different carbohydrate classes have remained unclear. Here, we identify an oral enzymatic-metabolic sensing axis involving maltase glucoamylase (MGAM) and glucokinase (GCK) that enables rapid detection of glucose-yielding saccharides. By locally preprocessing complex carbohydrates and coupling this activity to intracellular glucose sensing, the taste system can proactively bias intake toward energetically favorable foods prior to digestion. This mechanism provides a solution to the longstanding paradox of how glucose, a primary reinforcer of appetite, can shape ingestive behavior despite rarely being encountered in its free form in the diet^8,26^. Here, we identify an oral enzymatic-sensory axis involving maltase glucoamylase and glucokinase that enables the taste system to rapidly detect glucose-yielding saccharides. This mechanism facilitates the proactive targeted intake of metabolically valuable carbohydrates. We find that regular consumption of specific sugars induces expression of these enzymes in taste papillae, enhancing preference for saccharides that ultimately deliver higher glucose yields to the body. Disruption of either enzyme in major taste fields abolishes di- and poly-saccharide containing substances. Notably, while this pathway can function independently of canonical sweet receptors, it compensates for diminished sweet signaling to preserve oral sensitivity to dietary carbohydrates.

Sweet receptors are the main mechanism through which sugars engage ingestive behaviors. Polysaccharides engage a separate, yet to be identified receptor, but both signaling pathways utilize TRPM5 ^26,28^. Here, we found that mice deficient in TRPM5 are capable of tasting and responding in a positive fashion to monosaccharides and disaccharides, though at levels that were diminished relative to their TRPM5+ littermates, and B6 control mice. The robust concentration-dependent licking for maltose in TRPM5- was somewhat especially surprising given that mice that lack sweet receptors were previously found to be unresponsive to maltose and sucrose in brief access taste tests or taste detection tests^29,30^. However, we know from other studies that rodents can distinguish the taste of sucrose, the prototypical sweet sugar, from maltose and from even more complex carbohydrates, like Polycose^7,31,32,33^. Which has long suggested that alternative receptors exist. These data add to emerging lines of evidence that carbohydrates may engage other pathways, including gluco-specific sensors^14,15,17,34^. Importantly, we discovered that TRPM5- also had fewer Tas1r3 and *Gck* transcripts in their taste papillae, suggesting that taste signaling pathways may regulate one another, and this has significant implications for interpreting whether TRPM5 affects sensitivity via direct transduction or indirect regulatory mechanisms. In the context of the present studies, it is thus possible that we are underestimating the significance of TRPM5-independent signals (e.g., glucokinase) in basic sugar detection.

Dietary sugar conditions played an important role in tuning the relative salience of these nutritive taste signals. In particular, mice that were provided regular access to glucose and fructose, or sucrose and maltose, developed a special attraction to the taste of maltose. Maltose is generally considered less sweet than sucrose, but, once digested, yields more glucose than sucrose^35^. This hedonic shift was associated with an upregulation in glucokinase in the taste papilla. Acute and selective silencing of glucokinase in the major taste fields reverted these preferences in TRPM5+ and – mice, establishing that gustatory GCK is causally linked to the elevated acceptance of maltose.

Since maltose is not capable of activating GCK in its disaccharide form, we reasoned that an upstream mechanism capable of rapidly generating free glucose from maltose must also be involved. Sukumaran et al., found that disaccharide hydrolyzing enzymes (α-glucosidases) maltase-glucoamylase (MGAM) and sucrase-isomaltase (SIS), are expressed at the apical membrane of taste cells and interference with their activity using broad spectrum pharmacological blockers decreased gustatory nerve responses to sucrose and maltose, but whether these effects were relevant for taste-guided behavior was not assessed ^21^. Because the α-glucosidases that break down disaccharides are also present in the proximal gut and thus could be impacted by oral dosing of these anti-glycolytic compounds, we developed a shRNA-mediated virogenetic approach to silence MGAM activity in the major taste fields on the tongue, leaving all other sources intact, and allowing us to link lingual MGAM processing with taste-guided behaviors. In a verification of the approach, we found that MGAM KD reduced the size of licking bursts for maltose in a short consumption test in naïve mice. The licking burst size has been verified to increase linearly with sucrose concentration and burst number significantly decreases with food deprivation state^23,36^; thus, burst size can provide information of hedonic value that that is not revealed by volumetric intake alone^36^. Our findings provide the first evidence that MGAM activity around the taste cells contributes to the inherent appeal of this sugar. This supports the model that MGAM rapidly cleaves maltose and thus generates ligands in the form of free glucose for nearby glucosensors and sweet receptors to bolster carbohydrate detection^21,37^.

We discovered that lingual MGAM was elevated in mice that have diminished sweet sensing capabilities. TRPM5- mice were substantially less responsive to monosaccharides in the brief access taste test than B6 mice. These behavioral findings are in accord with the fact that TRPM5- mice had fewer transcripts for the sweet receptor gene (*Tas1r3*) and *Gck* in the taste buds. Notably, despite having impoverished machinery for sensing sweetness and simple sugar, TRPM5- had normal behavioral responsivity to maltose, an effect that was correlated with elevated levels of *Mgam* taste buds. This led us to suspect that MGAM may be adaptively upregulated to compensate for reduced sensitivity. This was corroborated by our findings in 129/J mice, which carry a *Tas1r3* polymorphism that renders them subsensitive to sweet. These mice licked as avidly for maltose as sweet-sensitive B6 and had significantly higher levels of *Mgam*. Several studies have already shown that 129/J display a greater preference for a cornstarch mixture compared to B6 mice and they have a preference for Polycose over water, confirming their avidity towards complex polysaccharides remains intact^38,39^.Our results reveal a novel mechanism whereby compensatory changes in upstream processing of oral saccharides preserves sensitivity to these nutrients by generating more ligands capable of binding to nearby receptors, including glucosensors.

Our results showed that both types of sugar exposure (glucose and fructose; maltose and sucrose) led to a significant increase in *Mgam* transcripts in the circumvallate papillae. Upregulated *Gck* and *Mgam* were associated with the increased avidity for glucose and maltose across studies. Interestingly, *Mgam* levels appear to be driven by the exposure to fructose content, not glucose. This differs from what has been observed in the gut^1^. It could be that in the absence of sufficient sources of glucose in the diet, *Mgam* is upregulated in a compensatory manner at the very first site of oral processing to promote the detection of foods that will ultimately yield more glucose. On the other hand, several studies have determined that excessive fructose or sucrose intake instigates metabolic disfunction^40,41,42,43^. Whether the diet-induced changes reflect an adaptive or maladaptive process remains to be seen.

All of the foods humans and other mammals normally consume comprise a mixture of macronutrients and common foods in the western diet are high in refined carbohydrates, including maltodextrins. Maltodextrins are readily cleaved by amylase, including in the saliva, to shorter glucose chains like maltose and maltotriose^44^. Thus, we assessed whether *Gck* and MGAM on the lingual epithelium play roles in consumption of a maltodextrin-containing mixed nutrient liquid diet, Peptamen, in a short licking test. Because all the mice were highly motivated to consume the diet in this task, we did not see overall differences in total licks (with a cap at 1000 licks) or other changes in the microstructural organization of licking behavior when averaged across the entire test session in mice that had reduced *Gck* or *Mgam* in the major taste fields. However, we did find that silencing either of these two lingual enzymes, *Gck* or *Mgam*, reduced the cumulative initial licking burst size relative to the control treatment on the post-KD test, suggesting that loss of either enzymatic mechanism impaired the initial detection and motivation to lick for the mixed nutrient shake. Given that refined carbohydrates engage other signaling pathways after they are ingested, this finding was not surprising but provides the first evidence, to our knowledge, that oral metabolic processing contributes to motivated consumption of a complex, mixed diet. More studies are needed now to examine the relative contribution of lingual *Gck* and *Mgam* to dietary intake, including with other nutrient compositions, and over the longer term.

### General Conclusion and Perspectives

Together, these findings reveal that the oral sensory system performs an active metabolic computation, rather than passively reporting chemical taste qualities. By coupling local enzymatic preprocessing of complex carbohydrates to intracellular glucose sensing, the taste epithelium can estimate the energetic value of food before ingestion and bias intake accordingly. This mechanism operates independently of canonical sweet taste receptors and is adaptively regulated by diet and sensory capacity, underscoring the flexibility of peripheral nutrient sensing. Given the prevalence of refined carbohydrates in modern diets, including maltodextrins that rapidly generate glucose oligomers in the mouth, oral metabolic sensing plays an important role in shaping early ingestive decisions, favoring more carbohydrate intake. More broadly, these results expand current models of taste function to include metabolic evaluation as a core component of sensory processing at the mouth–brain interface.

The findings of these studies solidify our previous work showing that rodents are tuned to detect glucose at the level of the taste system. Here we discovered that two enzymes, MGAM and GCK, work in tandem to break down then subsequently detect glucose molecules to reinforce consumption of complex carbohydrates. Loss of either mechanism significantly blunted the inherent and learned attraction to maltose, and the immediate attraction to a mixed nutrient diet containing maltodextrins. Like glucose, detection of maltose and by extension carbohydrates that break down to maltose do not appear to require the sweet- and/or canonical G-protein coupled receptor taste pathway. Further, digestive capacity at the taste cells via MGAM is upregulated to compensate for impoverished sweet reception, especially under dietary conditions associated with routine sugar intake. These findings contribute to the understanding of nutrient detection, food preferences and carbohydrate consuming behaviors. In human studies, GCK or MGAM could be tested to understand appetite regulation, sugar craving and individual variabilities to explain why some people may prefer high-sugar diets more than others. These glucosensors could become potential therapeutic targets through silencing or modulation as the metabolic response to sugars could be blunted to combat dietary diseases. As global rates of obesity and diabetes rise, understanding taste biology helps design better public health strategies.

## Methods

### Animals

Adult male B6 mice (n= 97) were purchased from Jackson Laboratories at approximately 9-10 weeks of age. TRPM5+ (♀:n=33; ♂:n=23) and TRPM5- (♀:n=21; ♂:n=15) mice were bred in-house from breeder pairs originally donated by Emily Liman (USC), T1RKO (♀:n=7; ♂:n=14) mice were bred in-house from breeder pairs originally donated by Alan Spector (Florida State University). TRPM5 + versus - status was determined behaviorally and genetically. Prior to any experiments, all mice from the TRPM5 colony were screened for their sensitivity to quinine (0.1mM) (a non-sweet, but TRPM5-dependent stimulus) in a 48-hour two bottle choice test (against plain water). A quinine: water discrimination score was calculated as total quinine intake divided by total water intake, with a score of 0.5 indicating no discrimination. At the conclusion of the experiments, we confirmed TRPM5 status by measuring *Trpm5* mRNA expression in the circumvallate papillae. Using the expression levels of the B6 mice as the standard, any mouse whose Trpm5 mRNA level was greater than the 5^th^ percentile of the B6 was TRPM5+; those below this threshold were confirmed TRPM5-. TRPM5+ mice were more sensitive to quinine (discrimination score = 0.284 ± 0.039) than TRPM5- mice (discrimination score: 0.443 ± 0.035).

Male 129/J (n=6) were purchased from Jackson Laboratories at approximately 10 weeks of age. All mice were acclimated to the experimental housing conditions for at least five days before the start of the study. The colony room was maintained on a 12:12 hour light/dark cycle, lights on at 6AM, with controlled temperature and humidity. All mice were individually housed in ventilated shoebox cages with a Nestlet for environmental enrichment. Pelleted chow (Pico Labdiet, #5053) was provided and water was available through an automated system (lixit), unless noted otherwise below for experimental purposes. All the experimental procedures were approved by *University of Southern California Institutional Animal Care and Use Committee*.

### Sugar Exposure

Approximately 24 hours before the start of this dietary intervention, chow was removed from the home cage, and mice were gradually reduced to 85% of their ad libitum body mass through a daily ration of chow and then maintained at this weight for the duration of the study. Once at 85% body weight, mice were randomly assigned to either the sugar exposure group or the sugar naïve group. The sugar exposure groups received a bottle containing 40 ml of a sugar solution each day. For the studies shown in **Figures 1, 2, 3, and 7**, 0.316 M glucose, 0.56 M glucose, 1.1 M glucose, 0.316 M fructose, 0.56 M fructose, and 1.1 M fructose were each provided for 24 hours in a randomized schedule, with each sugar concentration presented once per block over 3-5 successive blocks. In all of the aforementioned studies, sugar exposed mice had their lixit removed throughout the sugar exposure period, except the T1RKO mice, which had the lixit returned after the first block. For the study shown in **Figure 3H**, the glucose group received 0.316 M glucose, 0.56 M glucose, 1.1 M glucose only, the fructose group received 0.316 M fructose, 0.56 M fructose, and 1.1 M fructose only, and the glucose + fructose group received both sets of sugars with all groups on a randomized schedule. For the study shown in **Figure 4**, 40ml of 5% maltose, 10% maltose, 20% maltose, 5% sucrose, 10% sucrose, and 20% sucrose were provided instead. Due to the high mass concentration and viscosity of the disaccharides, we standardized those solutions by % w/v to closely match the monosaccharide mass load, thus better matching the daily caloric exposure of the glucose and fructose solutions. Intake was measured to the nearest gram. In all studies, a naïve group was given access to a water bottle throughout the exposure period. The sugar solutions were prepared fresh every day with deionized water (dH_2_O) and served at room temperature in addition to the chow ration. Three days following the sugar exposure period, sugar exposed and naïve mice were trained and tested for brief access taste tests in a gustometer or Med Associates Davis Rig^45,46^. Training and testing details outlined below.

### Brief Access Taste Testing

Mice were trained to lick for non-sweet fluids in the gustometer or Davis Rig similar to procedures outlined in Chometton et al., 2022. Briefly, water was removed from the home cage (at least 20 hours prior to the first training session), and then mice were trained to lick for at a stationary tube or sample ball for 20 minutes (dH_2_O or 4.5% corn oil emulsion with 0.6% sunflower lecithin; COE) for two sessions, run on consecutive days. In the gustometer, each lick was recorded by a load cell and activated a pump containing the training stimulus to dispense the fluid onto the sample ball (1 ul/lick). In the Davis Rig, the mice were trained to lick at a sipper tube through a controlled access slot in the testing chamber. In both apparatuses, registered licks were time-stamped for analyses. Water was returned to the home cage after the second training session. Next, mice were trained to lick for 4.5% COE in successive brief duration trials (10-s each, 7.5-s intertrial interval). Mice had to lick at least once to initiate a trial; once initiated the trial lasted 10-s irrespective of how many licks were made. Mice could initiate as many trials as possible in each 20-minute session and were required to achieve 14 trials within a 20-minute session (1-2 sessions) to proceed to testing. For brief access taste tests, the mice were presented with an array of stimulus solutions, presented in serial block randomized order without replacement. For mice in the studies shown in **Figures 1, 4 and 7**, the stimuli were dH_2_O, 0.316M, 0.56M, 1.1M of glucose and fructose. For the mice in studies shows in **Figures 2**, **3, and 7** the stimuli were dH_2_O, 0.316M, 0.56M, 1.1M of maltose and sucrose. For the study depicted in **Figure 4**, the stimuli were dH_2_O, 5%, 10% and 20% maltose and sucrose. For the study shown in **Figure 5**, the stimuli were dH_2_O, 0.075M, 0.15M, 0.3M, 0.6M and 1.0M maltose. The number of licks taken during each stimulus trial (10-s) was averaged across trials. A lick score was then calculated for each stimulus as follows: Lick score =average licks to the stimulus – average licks to dH_2_O.

### Single Solution Licking Tests

To measure lick avidity for a single stimulus, mice were first trained to lick in the gustometer as follows. First, water restricted mice were given 20-minute access to dH_2_O in the gustometer, where each lick delivered approximately 1µl of fluid; this was done for two sessions. Next, mice were given access to 0.3 M sucrose for 20-minute in the gustometer after an overnight fast. For maltose testing in **Figure 6**, mice were food deprived overnight and then given access to 0.6 M maltose until they took 300 licks or 10 minutes, whichever came sooner. For Peptamen testing in **Figure 8**, mice were food deprived overnight and then given access to unflavored Nestle Peptamen Jr (1 kcal/ml) for 1000 licks or 20 minutes, whichever came sooner. The microstructural organization of licking behaviors was analyzed in each test, according to the methods outlined in Chometton et al., 2022, where licking bursts were defined as runs of consecutive licks separated by < 1s.

### Collection of Taste Papillae

As described in Chometton et al., at the end of the study mice received an intraperitoneal (IP) injection with a lethal dose of Euthasol® (780 mg pentobarbital sodium and 100 mg phenytoin sodium per kg body weight)^10^. Mice were immediately decapitated and the whole tongue was extracted and pinned to a Sylgard® dish containing Tyrode’s buffer solution under the microscope. Then, 0.5ml of an enzyme cocktail (1.0 mg/ml collagenase/dispase A (11088793001, Sigma Aldrich)) in phosphate-buffered saline (PBS) was injected under the circumvallate papillae epithelium. The tongue was removed from the Sylgard® dish and placed into an oxygenated tube filled with 30-40ml of Tyrode’s buffer solution and gently agitated with oxygen at room temperature for 20min. Next, the whole tongue was again placed on the Sylgard® dish under the microscope and the circumvallate papillae epithelium was peeled and immediately placed in RNAlater (ThermoFisher Scientific) at 4°C overnight then placed in a -80°C freezer until processing with RT qPCR.

### Real-Time Quantitative PCR

Circumvallate papillae were assayed for mRNA expression levels of taste receptor type 1 member 3 (*Tas1r3*), glucokinase (*Gck*), maltase glucoamylase (*Mgam*), transient receptor potential cation channel subfamily M member 5 (*Trpm5*) and β-actin as the house keeping control gene. Total RNA was extracted from each tissue sample with RNeasy Micro Kit (#74004, Qiagen) following the manufacturer’s protocol. The RNA concentration was then measured using a NanoDrop Spectrophotometer (ND ONE-W, ThermoFisher Scientific). RNA samples underwent reverse transcription to be synthesized into cDNA using the QuantiTect Reverse Transcription Kit (#205311, Qiagen) according to the manufacturer’s protocol. Next, the cDNA samples were amplified for the genes of interest using the Taqman PreAmp Master Mix (#4391128, Applied Biosystems). Quantitative real-time PCR was then performed in triplicates on a 384-well plate with the TaqMan Fast Advanced Master Mix (#444557, Applied Biosystems) using the QuantStudio 5 Real-Time PCR System (Applied Biosystems). Non-template controls were included to verify the absence of genomic DNA contamination. The following TaqMan probes were used β-actin (Mm02619580_g1), *Gck* (Mm00439129_m1), *Mgam* (Mm01163791_m1), *Trpm5* (Mm01129032_m1) and *Tas1r3* (Mm00473459_g1). The reactions were run in triplicate and the results were normalized to β-actin expression. Expression of the control group was used to normalize the data. No template controls were used to verify absence of genomic DNA contamination. The ΔΔCt method was used to analyze the differences in expression levels of each gene between groups^47^.

### shRNA Knockdown Surgeries

Target genes of interest were silenced in the major taste fields according to our previously published protocol. Briefly, mice were anesthetized with isoflurane (5% induction rate; 2-3% maintenance rate, as needed) to receive a scrambled sequence shRNA (Control, sc-108080, Santa Cruz Biotechnology), GCK shRNA (sc-35459-V, Santa Cruz Biotechnology), or Maltase-Glucoamylase shRNA (m) (sc-75741-V, Santa Cruz Biotechnology). All shRNAs were packaged in a lentiviral vector. The virus was pipetted directly onto the fungiform and circumvallate papilla (2-3 µl/site).

### Statistical Analyses

The data are plotted with the mean and ± SEM in all graphs. Statistical analyses were preformed using the Prism software version 10.4.2 (GraphPad Software; CA, US) and are listed in Supplementary Tables (1-10). Student’s paired or unpaired t-test were performed for two group comparisons. Two-tailed t-tests were performed unless otherwise specified. In two-way ANOVAs the Brown-Forsythe correction was applied.

The p-value of p< 0.05 after correction were considered statistically significant. Mice lick score values were removed if they did not reach sufficient trial minimum (14 trials). All data sets were checked for outliers using the Grubbs’ test (alpha= 0.05); outliers were removed as indicated.

## Acknowledgements

All schematics created using BioRender®. This work was funded by the National Institutes of Health: R01DC018562 (to L.A.S.) and F31DC021376 (to A.S.R.)

---

**Supplementary Fig. 1:**
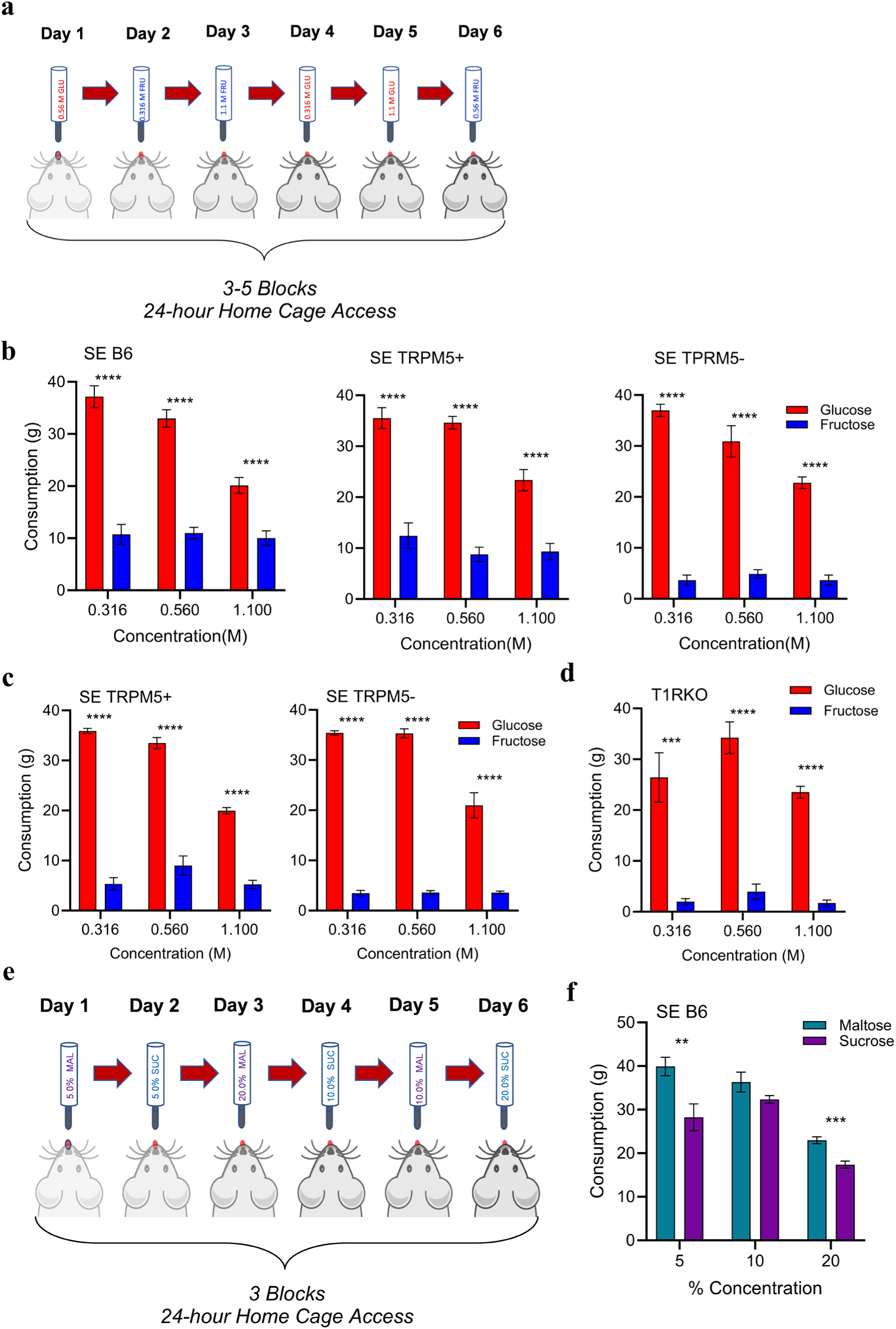
Sugar Exposure Paradigm. **a** Schematic depicting sugar exposure paradigm with glucose (0.316M, 0.56M and 1.1M) and fructose (0.316M, 0.56M and 1.1M) home cage access. **b** Mean (±SEM) glucose and fructose consumption during the last block (5 blocks; 30 days) for sugar exposed B6, TRPM5+ and TRPM5- (n=10/group) from Figure 1. **c** Mean (±SEM) consumption of glucose and fructose during the last block (3 blocks;18 days) TRPM5+ (n=7) and TRPM5- (n=9) from Figure 3. **d** Mean (±SEM) consumption of glucose and fructose for T1RKO (n=12) during the last block (3 blocks; 18 days) of sugar exposure from Figure 7. **e** Schematic depicting sugar exposure paradigm with maltose (5.0%, 10.0%, 20.0%) and sucrose (5.0%, 10.0%, 20.0%) home cage access. **f** Mean (±SEM) consumption of maltose and sucrose during last block (3 blocks; 18 days) of sugar exposure (SE) in B6 mice (n=8) from Figure 4. (*:p<0.05, **:p<0.01, ***:p<0.001, ****:p<0.0001). Statistical Analysis are in Supplementary Table 9.

**Supplementary Fig. 2:**
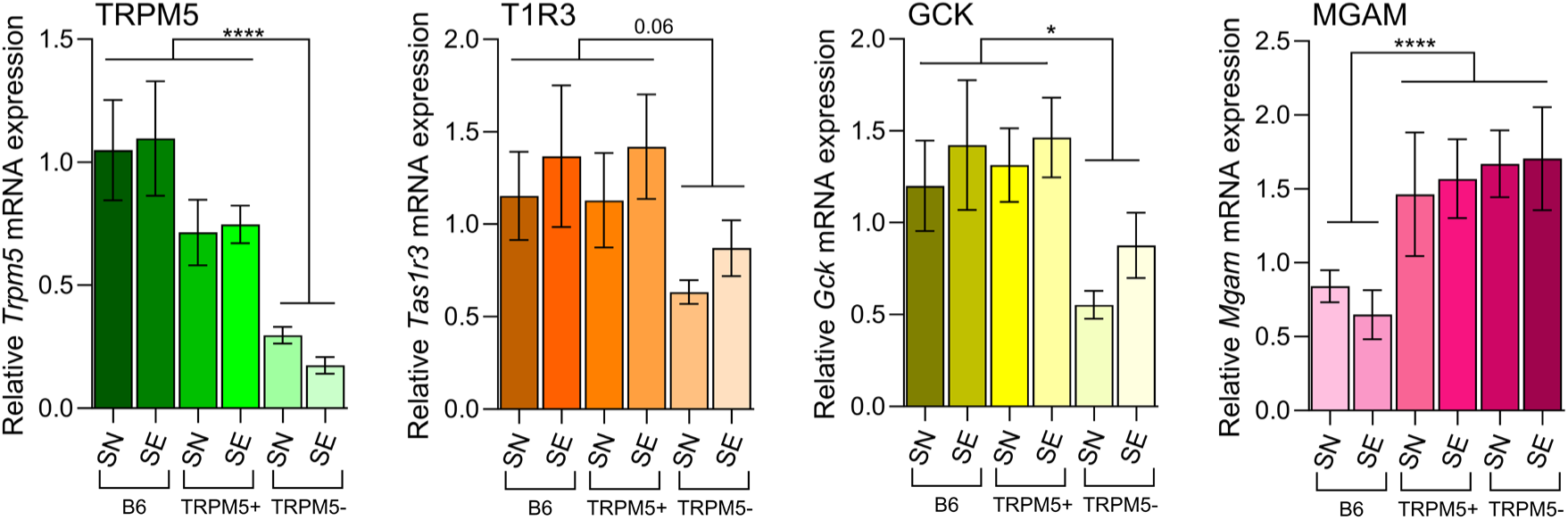
Differing taste gene expression in sugar exposed and sugar naïve sweet sensitive and subsensitive mice. Genetic expression for animals in Figure 1b and Figure 3g separated by SN or SE condition. Mean (±SEM) relative transcript expression of Trpm5 for B6 (SN: n= 8, SE: n= 8), TRPM5+ (SN: n= 5, SE: n= 6), TRPM5-(SN: n= 4, SE: n= 9); Tas1r3 for B6 (SN: n= 8, SE: n= 8), TRPM5+ (SN: n= 5, SE: n= 6), TRPM5- (SN: n= 4, SE: n= 9); Gck for B6 (SN: n= 8, SE: n= 8), TRPM5+ (SN: n= 5, SE: n= 5), TRPM5- (SN: n= 4, SE: n= 9); Mgam for B6 (SN: n= 7, SE: n= 5), TRPM5+ (SN: n= 4, SE: n= 5), TRPM5- (SN: n= 4, SE: n= 5). (*:p<0.05,****:p<0.0001). Statistical Analysis are in Supplementary Table 10.

**Supplementary Fig. 3:**
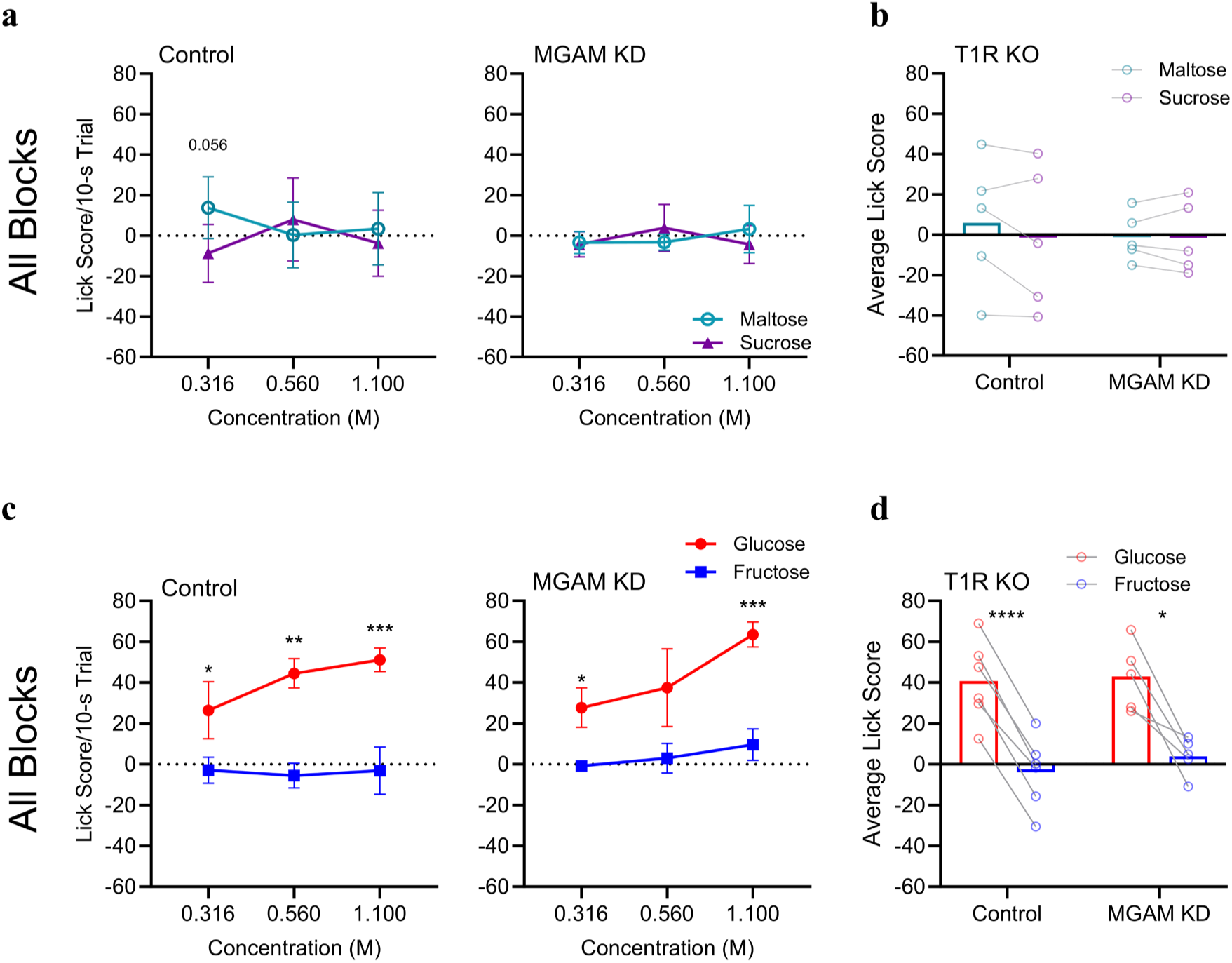
Sweet receptor knockout mice lick scores for all blocks after lingual silencing of MGAM. **a** Mean (±SEM) lick scores for all blocks of 0.316M, 0.56M, 1.1M maltose and sucrose in control (n=5) and MGAM KD (n=5) T1RKO. **b** Mean lick scores averaged across concentration for control and MGAM KD (5/group). **c** All blocks mean (±SEM) lick scores for 0.316M, 0.56M, 1.1M glucose and fructose in control (n=6) and MGAM KD (n=5) T1RKO. **d** Mean glucose and fructose lick scores averaged across concentration for control (n=6) and MGAM KD (n=5) T1RKO. (*:p<0.05, **:p<0.01, ***:p<0.001, ****:p<0.0001). Complementary data in Figure 7. Statistical Analysis are in Supplementary Table 11.

**Supplementary Table 1A:**
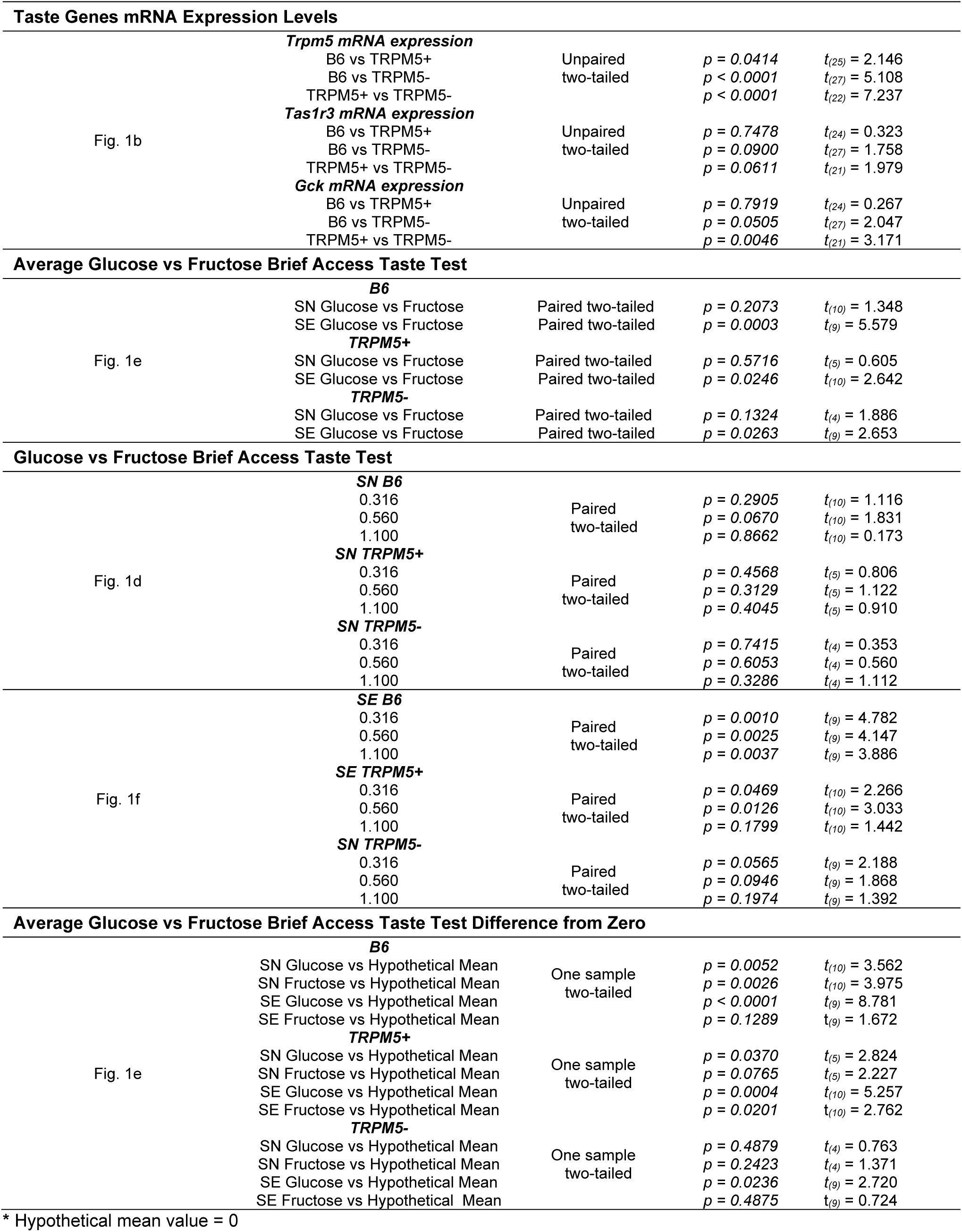
Student’s *t*-tests.

**Supplementary Table 1B:**
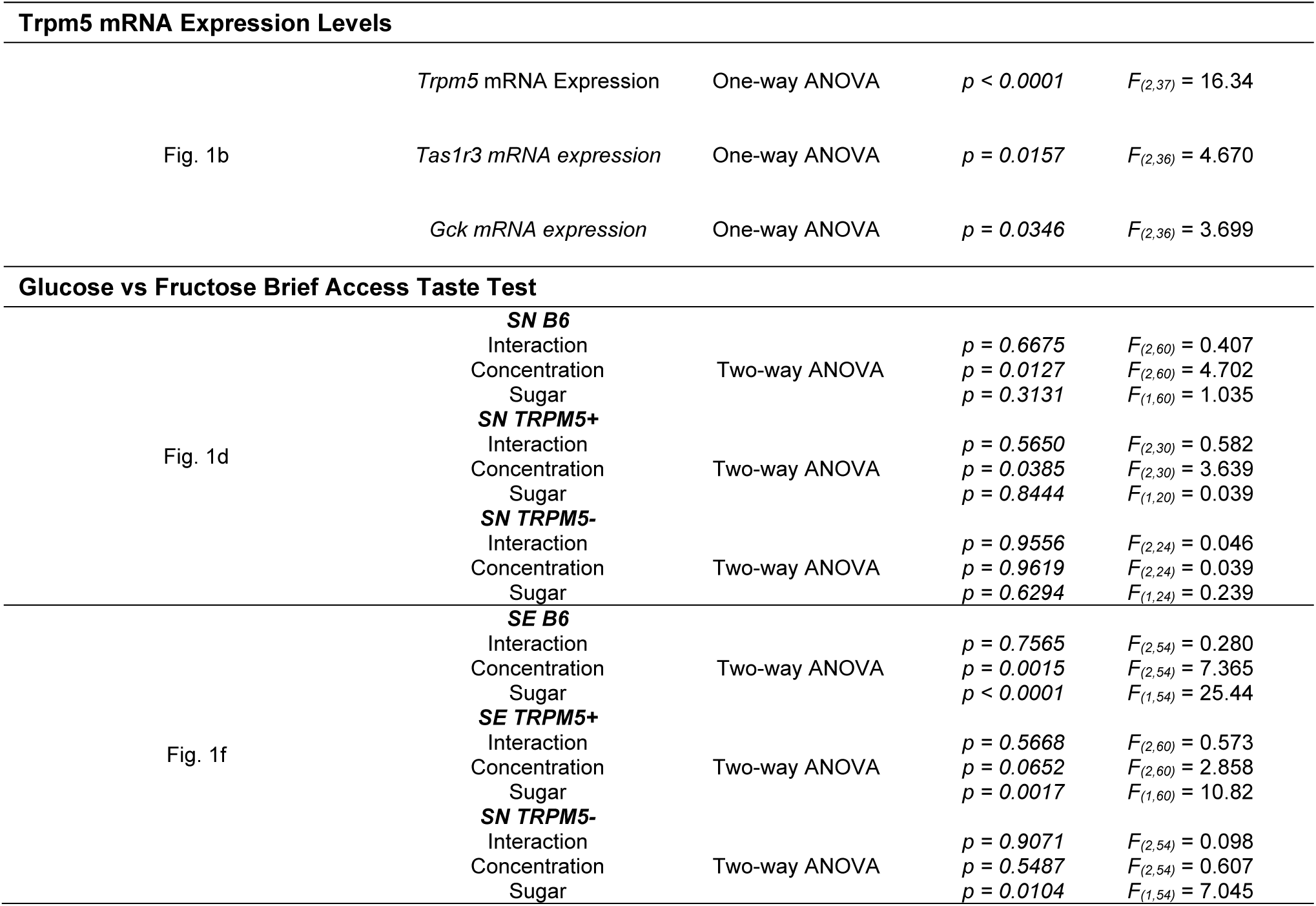
ANOVA tests.

**Supplementary Table 2A:**
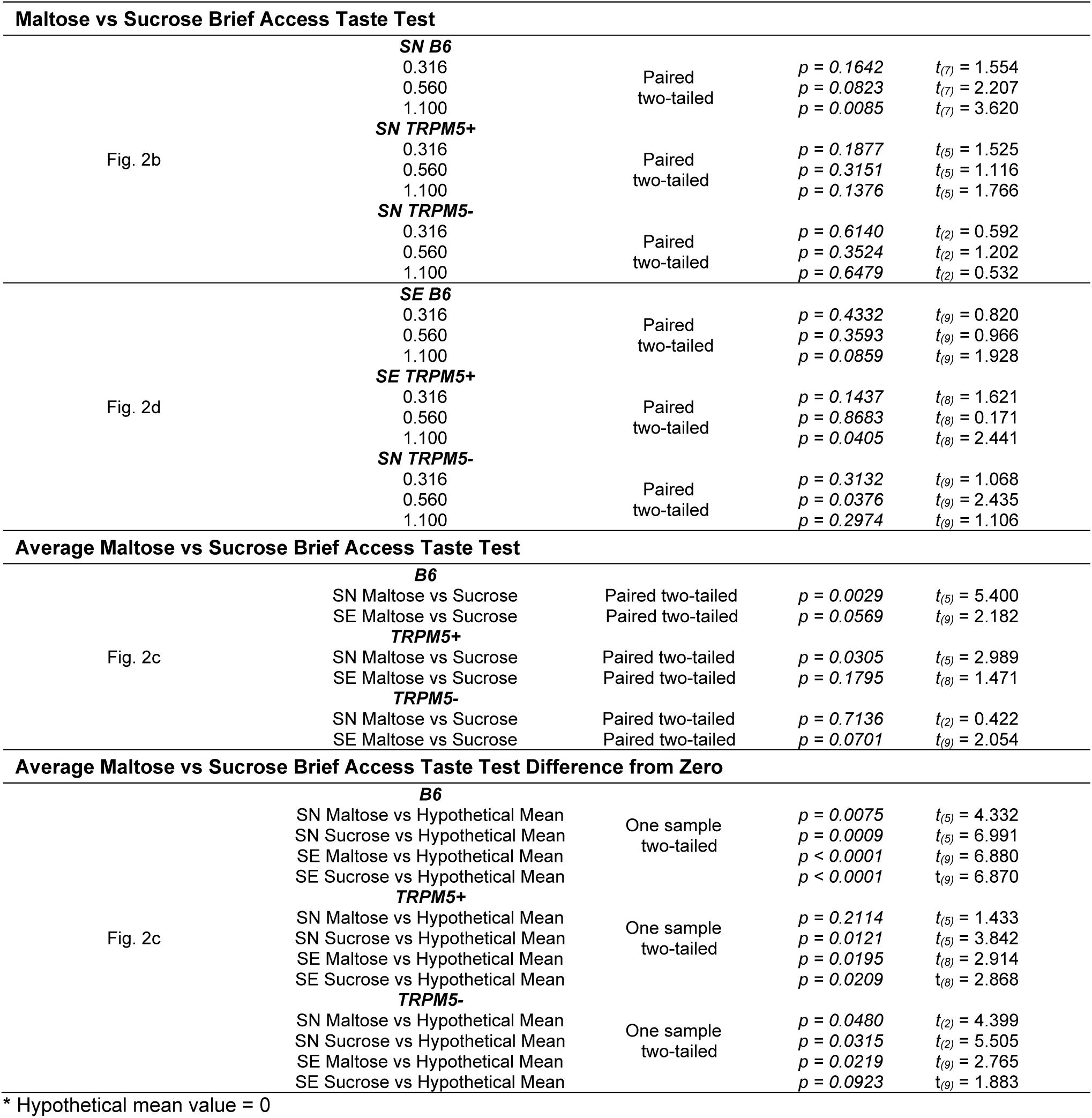
Student’s *t*-tests.

**Supplementary Table 2B:**
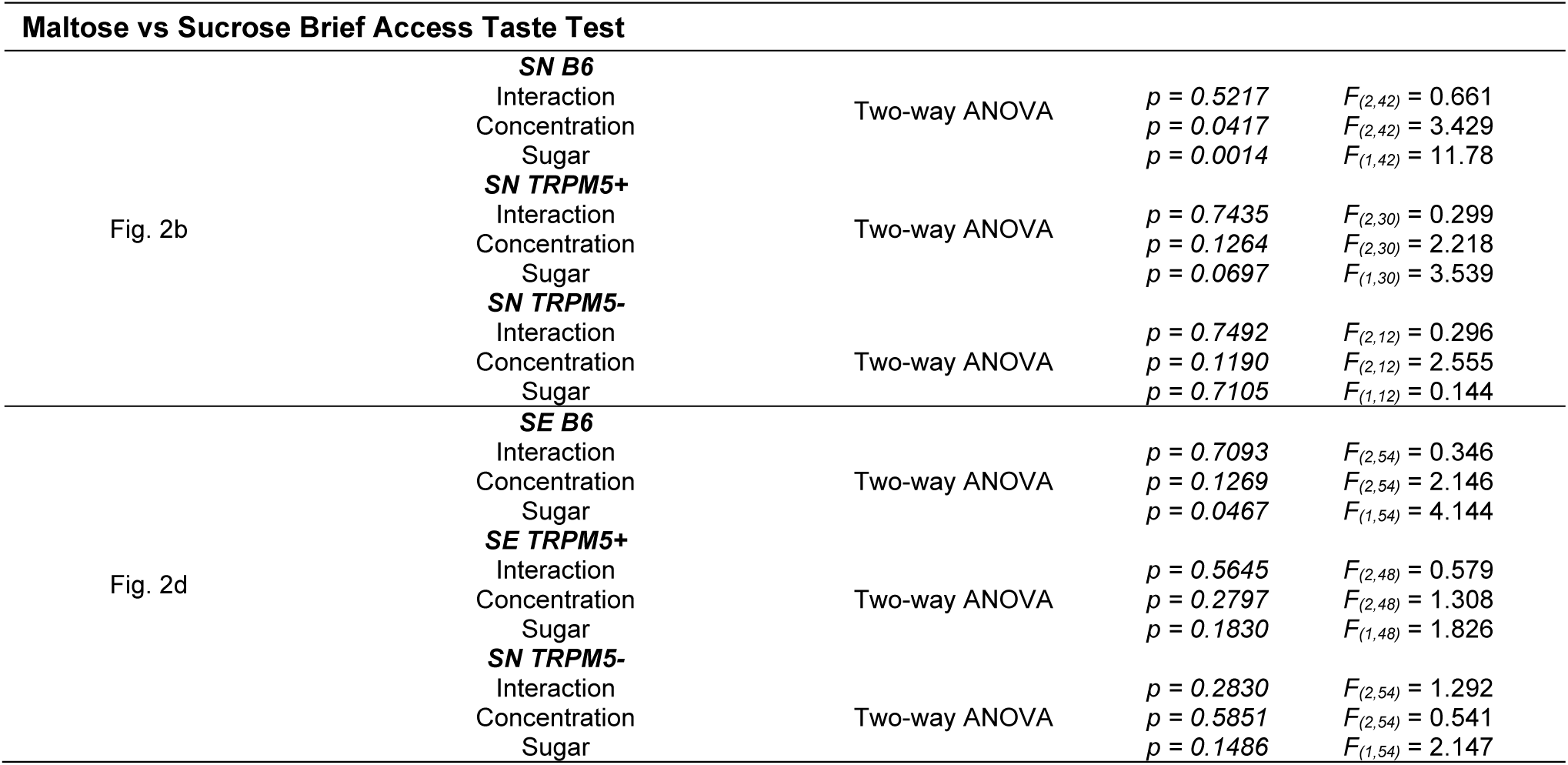
ANOVA tests.

**Supplementary Table 3A:** Student’s *t*-tests.

| TRPM5+ Maltose vs Sucrose Brief Access Taste Test |  |  |  |  |
| --- | --- | --- | --- | --- |
| Fig. 3c | *Control* |  |  |  |
| | 0.316 | Paired<br>two-tailed | $p = 0.4350$ | $t_{(10)} = 0.813$ |
| | 0.560 | | $p = 0.0252$ | $t_{(10)} = 2.628$ |
| | 1.100 | | $p = 0.6221$ | $t_{(10)} = 0.509$ |
|  | *GCK KD* |  |  |  |
| | 0.316 | Paired<br>two-tailed | $p = 0.0010$ | $t_{(11)} = 4.437$ |
| | 0.560 | | $p = 0.0203$ | $t_{(11)} = 2.710$ |
| | 1.100 | | $p = 0.7255$ | $t_{(11)} = 0.360$ |
| TRPM5+ Average Maltose vs Sucrose Brief Access Taste Test |  |  |  |  |
| Fig. 3d | Control Maltose vs Sucrose | Paired two-tailed | $p = 0.3767$ | $t_{(10)} = 0.925$ |
| | GCK KD Maltose vs Sucrose | Paired two-tailed | $p = 0.0041$ | $t_{(11)} = 3.614$ |
| | Control Maltose vs GCK KD Maltose | Unpaired two-tailed | $p = 0.1132$ | $t_{(21)} = 1.653$ |
| | Control Sucrose vs GCK KD Sucrose | Unpaired two-tailed | $p = 0.9136$ | $t_{(21)} = 0.110$ |
| TRPM5- Maltose vs Sucrose Brief Access Taste Test |  |  |  |  |
| Fig. 3e | *Control* |  |  |  |
| | 0.316 | Paired<br>two-tailed | $p = 0.0403$ | $t_{(9)} = 2.394$ |
| | 0.560 | | $p = 0.0063$ | $t_{(9)} = 3.543$ |
| | 1.100 | | $p = 0.9376$ | $t_{(9)} = 0.081$ |
|  | *GCK KD* |  |  |  |
| | 0.316 | Paired<br>two-tailed | $p = 0.3958$ | $t_{(10)} = 0.887$ |
| | 0.560 | | $p = 0.9435$ | $t_{(10)} = 0.073$ |
| | 1.100 | | $p = 0.4263$ | $t_{(10)} = 0.829$ |
| TRPM5- Average Maltose vs Sucrose Brief Access Taste Test |  |  |  |  |
| Fig. 3f | Control Maltose vs Sucrose | Paired two-tailed | $p = 0.0106$ | $t_{(9)} = 3.212$ |
| | GCK KD Maltose vs Sucrose | Paired two-tailed | $p = 0.3411$ | $t_{(10)} = 0.999$ |
| | Control Maltose vs GCK KD Maltose | Unpaired two-tailed | $p = 0.2122$ | $t_{(19)} = 1.291$ |
| | Control Sucrose vs GCK KD Sucrose | Unpaired two-tailed | $p = 0.6280$ | $t_{(19)} = 0.493$ |
| Mgam mRNA Expression Levels |  |  |  |  |
| Fig. 3g | *Mgam mRNA expression* |  |  |  |
| | B6 vs TRPM5+ | Unpaired<br>two-tailed | $p = 0.0066$ | $t_{(18)} = 3.070$ |
| | B6 vs TRPM5- | | $p = 0.0003$ | $t_{(19)} = 4.483$ |
| | TRPM5+ vs TRPM5- | | $p = 0.4306$ | $t_{(15)} = 0.810$ |
| Differing Exposure Mgam mRNA Expression Levels |  |  |  |  |
| Fig. 3h | *Mgam mRNA expression* |  |  |  |
| | Naive vs Hypothetical Mean* | One sample<br>one-tailed | $p = 0.4706$ | $t_{(1)} = 0.093$ |
| | Glucose vs Hypothetical Mean* | | $p = 0.3417$ | $t_{(4)} = 0.439$ |
| | Fructose vs Hypothetical Mean* | | $p = 0.0440$ | $t_{(5)} = 2.115$ |
| | Glucose+Fructose vs Hypothetical Mean* | | $p = 0.0515$ | $t_{(4)} = 2.106$ |

\*Hypothetical mean value = 1

**Supplementary Table 3B:** ANOVA tests.

| TRPM5+ Maltose vs Sucrose Brief Access Taste Test |  |  |  |  |
| --- | --- | --- | --- | --- |
| Fig. 3c | **Control** |  |  |  |
| | Interaction | | $p = 0.1060$ | $F_{(2,60)} = 2.331$ |
| | Concentration | Two-way ANOVA | $p < 0.0001$ | $F_{(2,60)} = 14.47$ |
| | Sugar | | $p = 0.2569$ | $F_{(1,60)} = 1.310$ |
|  | **GCK KD** |  |  |  |
| | Interaction | | $p = 0.2871$ | $F_{(2,66)} = 1.272$ |
| | Concentration | Two-way ANOVA | $p = 0.0298$ | $F_{(2,66)} = 3.705$ |
| | Sugar | | $p = 0.0064$ | $F_{(1,66)} = 7.925$ |
| TRPM5+ Average Maltose vs Sucrose Brief Access Taste Test |  |  |  |  |
| Fig. 3d | Interaction | | $p = 0.2144$ | $F_{(1,42)} = 1.589$ |
| | Group (Con or GCK KD) | Two-way ANOVA | $p = 0.2745$ | $F_{(1,42)} = 1.226$ |
| | Sugar | | $p = 0.0287$ | $F_{(1,42)} = 5.133$ |
| TRPM5- Maltose vs Sucrose Brief Access Taste Test |  |  |  |  |
| Fig. 3e | **Control** |  |  |  |
| | Interaction | | $p = 0.3594$ | $F_{(2,54)} = 1.043$ |
| | Concentration | Two-way ANOVA | $p = 0.3249$ | $F_{(2,54)} = 1.148$ |
| | Sugar | | $p = 0.0577$ | $F_{(1,54)} = 3.759$ |
|  | **GCK KD** |  |  |  |
| | Interaction | | $p = 0.9534$ | $F_{(2,60)} = 0.048$ |
| | Concentration | Two-way ANOVA | $p = 0.9559$ | $F_{(2,60)} = 0.045$ |
| | Sugar | | $p = 0.6238$ | $F_{(1,60)} = 0.243$ |
| TRPM5- Average Maltose vs Sucrose Brief Access Taste Test |  |  |  |  |
| Fig. 3f | Interaction | | $p = 0.5431$ | $F_{(1,38)} = 0.377$ |
| | Group (Con or GCK KD) | Two-way ANOVA | $p = 0.2075$ | $F_{(1,38)} = 1.644$ |
| | Sugar | | $p = 0.2866$ | $F_{(1,38)} = 1.168$ |
| Mgam Genes mRNA Expression Levels |  |  |  |  |
| Fig. 3g | *Mgam* mRNA expression | One-way ANOVA | $p = 0.0012$ | $F_{(2,26)} = 8.755$ |
| Differing Exposure Mgam mRNA Expression Levels |  |  |  |  |
| Fig. 3h | *Mgam* mRNA Expression | One-way ANOVA | $p = 0.1695$ | $F_{(3,14)} = 1.940$ |

**Supplementary Table 4A:** Student’s *t*-tests.

| Maltose vs Sucrose Brief Access Taste Test |  |  |  |  |
| --- | --- | --- | --- | --- |
| Fig. 4b | **Sugar Naive** |  |  |  |
| | 5.00% | Paired<br>two-tailed | $p = 0.0218$ | $t_{(7)} = 2.938$ |
| | 10.0% | | $p = 0.0002$ | $t_{(7)} = 7.330$ |
| | 20.0% | | $p < 0.0001$ | $t_{(7)} = 12.51$ |
|  | **Sugar Exposed** |  |  |  |
| | 5.00% | Paired<br>two-tailed | $p = 0.1971$ | $t_{(7)} = 1.425$ |
| | 10.0% | | $p = 0.2410$ | $t_{(7)} = 1.281$ |
| | 20.0% | | $p = 0.7651$ | $t_{(7)} = 0.311$ |
| Average Maltose vs Sucrose Brief Access Taste Test |  |  |  |  |
| Fig. 4c | SN Maltose vs Sucrose | Paired two-tailed | $p < 0.0001$ | $t_{(7)} = 13.56$ |
| | SE Maltose vs Sucrose | Paired two-tailed | $p = 0.2687$ | $t_{(7)} = 1.201$ |
| | SN Maltose vs SE Maltose | Unpaired two-tailed | $p = 0.0038$ | $t_{(14)} = 3.469$ |
| | SN Sucrose vs SE Sucrose | Unpaired two-tailed | $p = 0.0460$ | $t_{(14)} = 2.189$ |
| Glucose vs Fructose Brief Access Taste Test |  |  |  |  |
| Fig. 4e | **Sugar Naive** |  |  |  |
| | 0.316 | Paired<br>two-tailed | $p = 0.3283$ | $t_{(7)} = 1.051$ |
| | 0.560 | | $p = 0.0331$ | $t_{(7)} = 2.646$ |
| | 1.100 | | $p = 0.8150$ | $t_{(7)} = 0.243$ |
|  | **Sugar Exposed** |  |  |  |
| | 0.316 | Paired<br>two-tailed | $p = 0.9051$ | $t_{(7)} = 1.237$ |
| | 0.560 | | $p = 0.0455$ | $t_{(7)} = 2.428$ |
| | 1.100 | | $p = 0.2421$ | $t_{(7)} = 1.278$ |
| Average Glucose vs Fructose Brief Access Taste Test |  |  |  |  |
| Fig. 4f | SN Glucose vs Fructose | Paired two-tailed | $p = 0.1620$ | $t_{(7)} = 1.563$ |
| | SE Glucose vs Fructose | Paired two-tailed | $p = 0.9094$ | $t_{(7)} = 0.118$ |
| | SN Glucose vs SE Glucose | Unpaired two-tailed | $p = 0.5674$ | $t_{(14)} = 0.586$ |
| Taste Genes mRNA Expression Levels |  |  |  |  |
| Fig. 4g | **Tas1R3 mRNA expression** |  |  |  |
| | Sugar Naive vs Sugar Exposed | Unpaired<br>two-tailed | $p = 0.4991$ | $t_{(14)} = 0.694$ |
|  | **GCK mRNA expression** |  |  |  |
| | Sugar Naive vs Sugar Exposed | Unpaired<br>two-tailed | $p = 0.0369$ | $t_{(14)} = 2.571$ |
|  | **Mgam mRNA expression** |  |  |  |
| | Sugar Naive vs Sugar Exposed | Unpaired<br>two-tailed | $p = 0.0091$ | $t_{(3)} = 6.049$ |

**Supplementary Table 4B:**
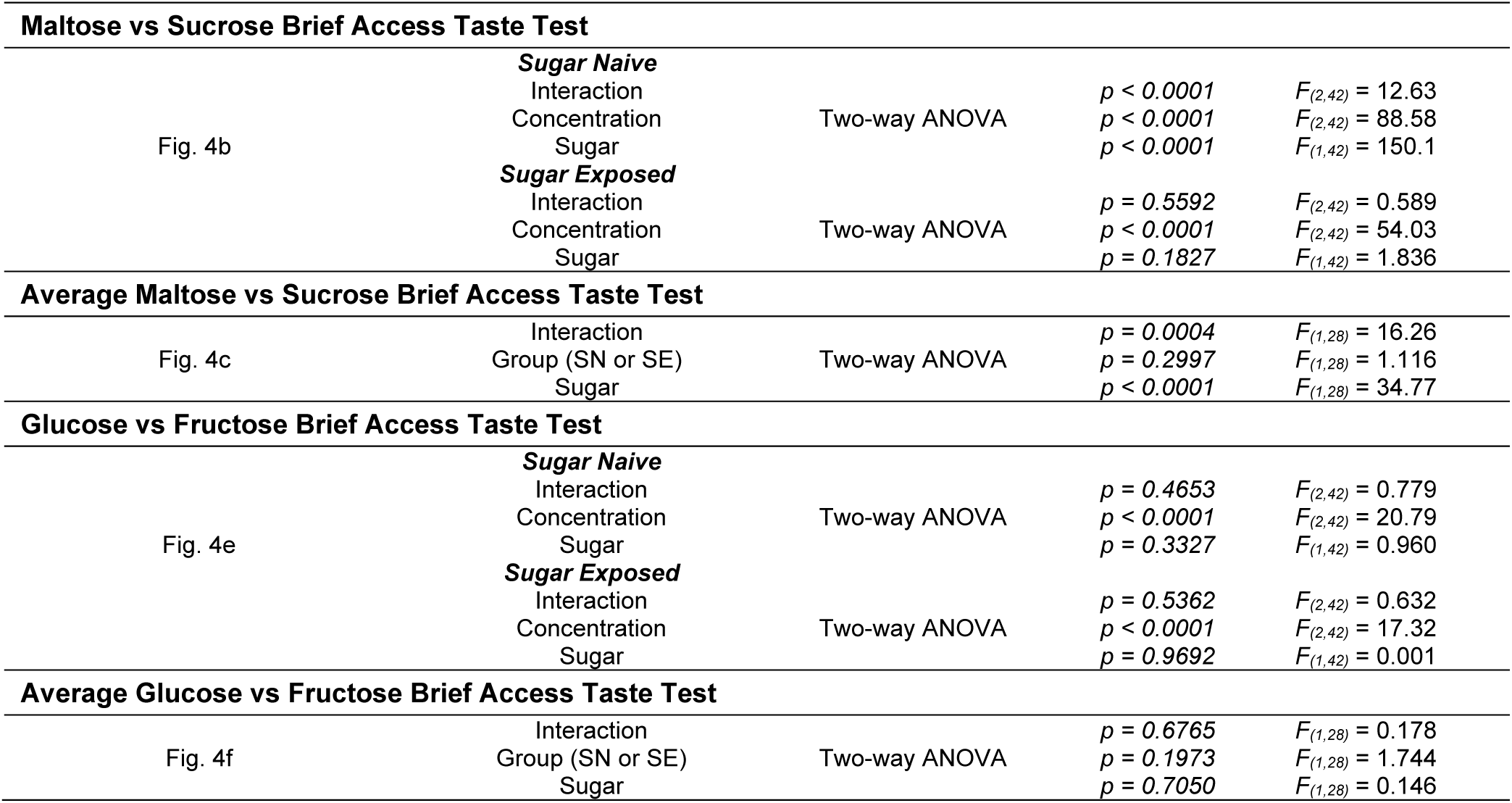
ANOVA tests.

**Supplementary Table 5A:** Student’s *t*-tests.

| Taste Genes mRNA Expression Levels |  |  |  |  |
| --- | --- | --- | --- | --- |
| Fig. 5b | **Tas1R3 mRNA expression** |  |  |  |
| | B6 vs 129 | Unpaired two-tailed | $p = 0.0033$ | $t_{(12)} = 3.655$ |
|  | **Gck mRNA expression** |  |  |  |
| | B6 vs 129 | Unpaired two-tailed | $p = 0.5279$ | $t_{(12)} = 0.650$ |
|  | **Mgam mRNA expression** |  |  |  |
| | B6 vs 129 | Unpaired two-tailed | $p = 0.0117$ | $t_{(12)} = 2.971$ |
| Maltose Brief Access Taste Test |  |  |  |  |
| Fig. 5d | **B6 vs 129** |  |  |  |
| | 0.075 | Unpaired two-tailed | $p = 0.8116$ | $t_{(4)} = 0.255$ |
| | 0.150 | | $p = 0.5169$ | $t_{(4)} = 0.710$ |
| | 0.300 | | $p = 0.8266$ | $t_{(4)} = 0.234$ |
| | 0.600 | | $p = 0.1485$ | $t_{(4)} = 1.787$ |
| | 1.000 | | $p = 0.4275$ | $t_{(4)} = 0.882$ |

**Supplementary Table 5B:** ANOVA tests.

| Maltose Brief Access Taste Test |  |  |  |  |
| --- | --- | --- | --- | --- |
| Fig. 5d | Interaction<br>Concentration<br>Group (B6 or 129) | Two-way ANOVA | $p = 0.6439$<br>$p = 0.0071$<br>$p = 0.0830$ | $F_{(4,20)} = 0.634$<br>$F_{(4,20)} = 4.788$<br>$F_{(1,20)} = 3.330$ |

**Supplementary Table 6:** Student’s *t*-tests.

| Taste Genes mRNA Expression Levels |  |  |  |  |
| --- | --- | --- | --- | --- |
| Fig. 6b | ***Mgam* mRNA expression** |  |  |  |
| | Control vs Hypothetical mean* | One sample | $p = 0.3856$ | $t_{(4)} = 0.311$ |
| | MGAM KD vs Hypothetical mean* | one tailed | $p = 0.0578$ | $t_{(3)} = 2.195$ |
|  | ***Gck* mRNA expression** |  |  |  |
| | Control vs MGAM KD | Unpaired<br>two-tailed | $p = 0.1062$ | $t_{(7)} = 1.854$ |
|  | ***Tas1R3* mRNA expression** |  |  |  |
| | Control vs MGAM KD | Unpaired<br>two-tailed | $p = 0.8042$ | $t_{(7)} = 0.257$ |
| Control vs MGAM KD 300 Lick Test |  |  |  |  |
| Fig. 6f | Burst Size | Unpaired | $p = 0.0259$ | $t_{(4)} = 3.456$ |
| | Burst Number | two-tailed | $p = 0.0437$ | $t_{(4)} = 2.910$ |
| | Total Licks | | $p > 0.9999$ | $t_{(4)} = 0.000$ |

\* Hypothetical mean value = 1

**Supplementary Table 7A:** Student’s *t*-tests.

| Taste Genes mRNA Expression Levels |  |  |  |  |
| --- | --- | --- | --- | --- |
| Fig. 7b | *Gck* mRNA expression |  |  |  |
| | Naive vs Hypothetical mean* | One sample | $p = 0.7893$ | $t_{(3)} = 0.292$ |
| | Sugar Exp vs Hypothetical mean* | one-tailed | $p = 0.0042$ | $t_{(3)} = 7.942$ |
|  | *Mgam* mRNA expression |  |  |  |
| | Naive vs Hypothetical mean* | One sample | $p = 0.7036$ | $t_{(3)} = 0.419$ |
| | Sugar Exp vs Hypothetical mean* | one-tailed | $p = 0.0490$ | $t_{(3)} = 2.376$ |
| Block 1 Maltose vs Sucrose Brief Access Taste Test |  |  |  |  |
| Fig. 7e | *Control* |  |  |  |
| | 0.316 | Paired<br>two-tailed | $p = 0.0568$ | $t_{(4)} = 2.653$ |
| | 0.560 | | $p = 0.5046$ | $t_{(4)} = 0.732$ |
| | 1.100 | | $p = 0.2128$ | $t_{(4)} = 1.481$ |
|  | *MGAM KD* |  |  |  |
| | 0.316 | Paired<br>two-tailed | $p = 0.1292$ | $t_{(4)} = 1.907$ |
| | 0.560 | | $p = 0.5241$ | $t_{(4)} = 0.697$ |
| | 1.100 | | $p = 0.2417$ | $t_{(4)} = 1.373$ |
| Block 1 Average Maltose vs Sucrose Brief Access Taste Test |  |  |  |  |
| Fig. 7f | Control Maltose vs Sucrose | Paired two-tailed | $p = 0.0513$ | $t_{(4)} = 2.752$ |
| | MGAM KD Maltose vs Sucrose | Paired two-tailed | $p = 0.2263$ | $t_{(4)} = 1.429$ |
| | Control Maltose vs MGAM KD Maltose | Unpaired two-tailed | $p = 0.2364$ | $t_{(8)} = 1.280$ |
| | Control Sucrose vs MGAM KD Sucrose | Unpaired two-tailed | $p = 0.6256$ | $t_{(8)} = 0.507$ |
| Block 1 Glucose vs Fructose Brief Access Taste Test |  |  |  |  |
| Fig. 7h | *Control* |  |  |  |
| | 0.316 | Paired<br>two-tailed | $p = 0.0376$ | $t_{(5)} = 2.809$ |
| | 0.560 | | $p = 0.1051$ | $t_{(5)} = 1.976$ |
| | 1.100 | | $p = 0.0212$ | $t_{(5)} = 3.310$ |
|  | *MGAM KD* |  |  |  |
| | 0.316 | Paired<br>two-tailed | $p = 0.0286$ | $t_{(4)} = 3.350$ |
| | 0.560 | | $p = 0.3426$ | $t_{(4)} = 1.076$ |
| | 1.100 | | $p = 0.0254$ | $t_{(4)} = 3.478$ |
| Block 1 Average Glucose vs Fructose Brief Access Taste Test |  |  |  |  |
| Fig. 7i | Control Glucose vs Fructose | Paired two-tailed | $p = 0.0033$ | $t_{(5)} = 5.245$ |
| | MGAM KD Glucose vs Fructose | Paired two-tailed | $p = 0.0160$ | $t_{(4)} = 4.010$ |
| | Control Glucose vs MGAM KD Glucose | Unpaired two-tailed | $p = 0.8423$ | $t_{(9)} = 0.205$ |

\* Hypothetical mean value = 1

**Supplementary Table 7B:**
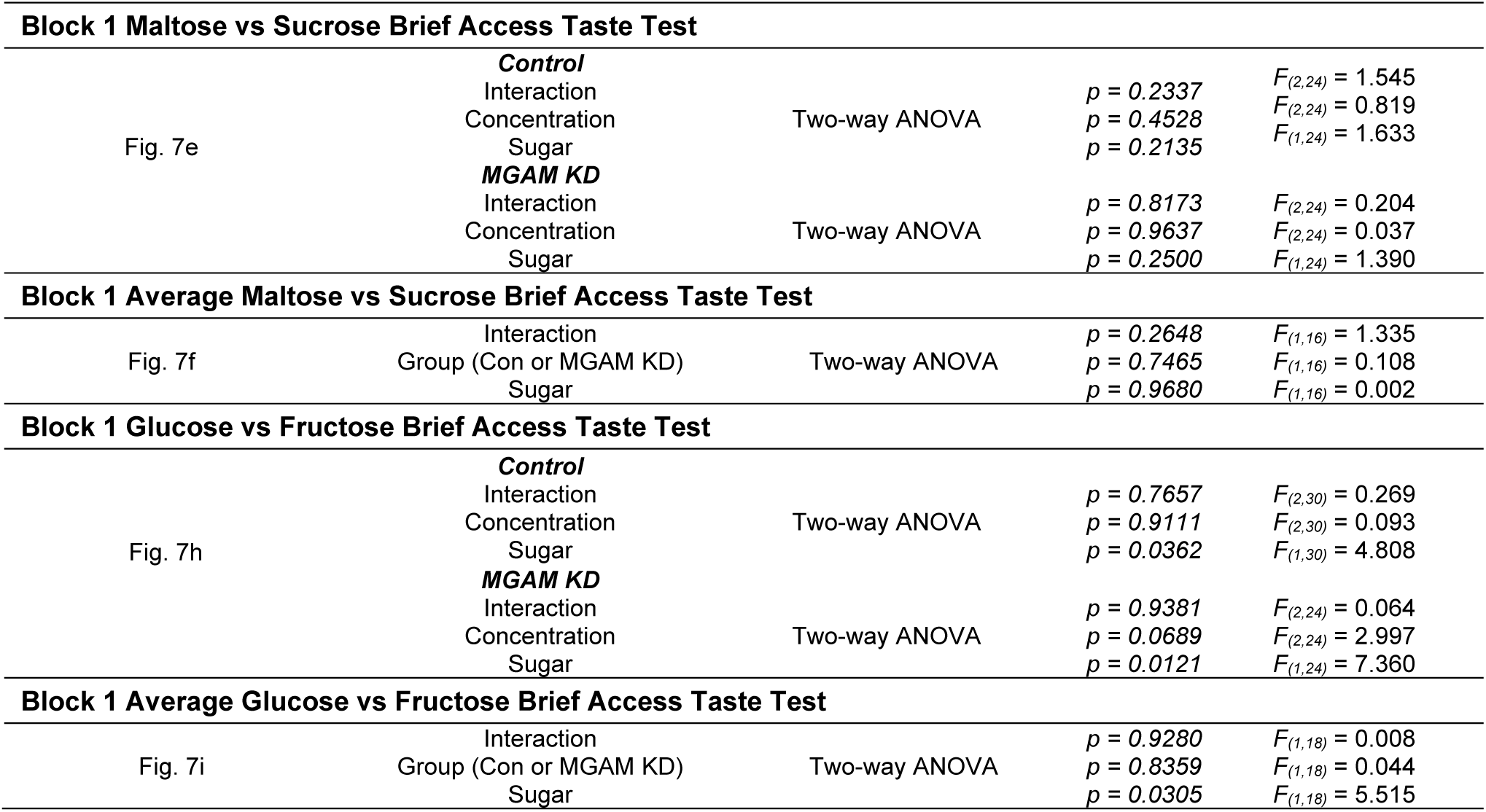
ANOVA tests.

**Supplementary Table 8A:**
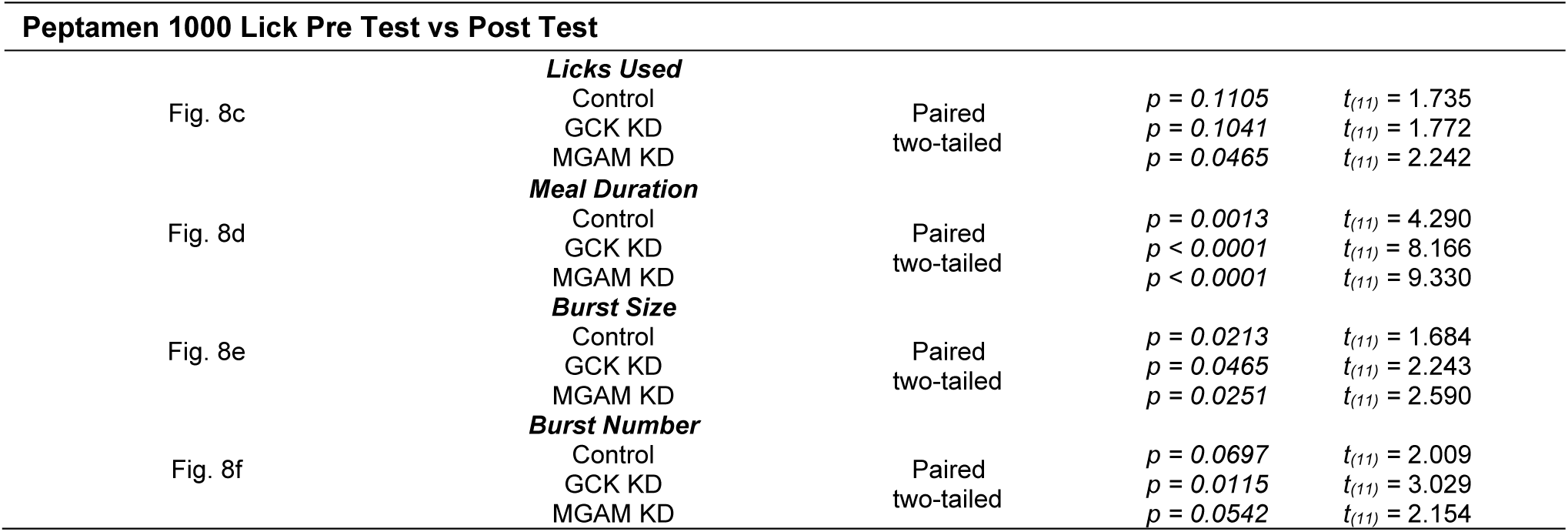
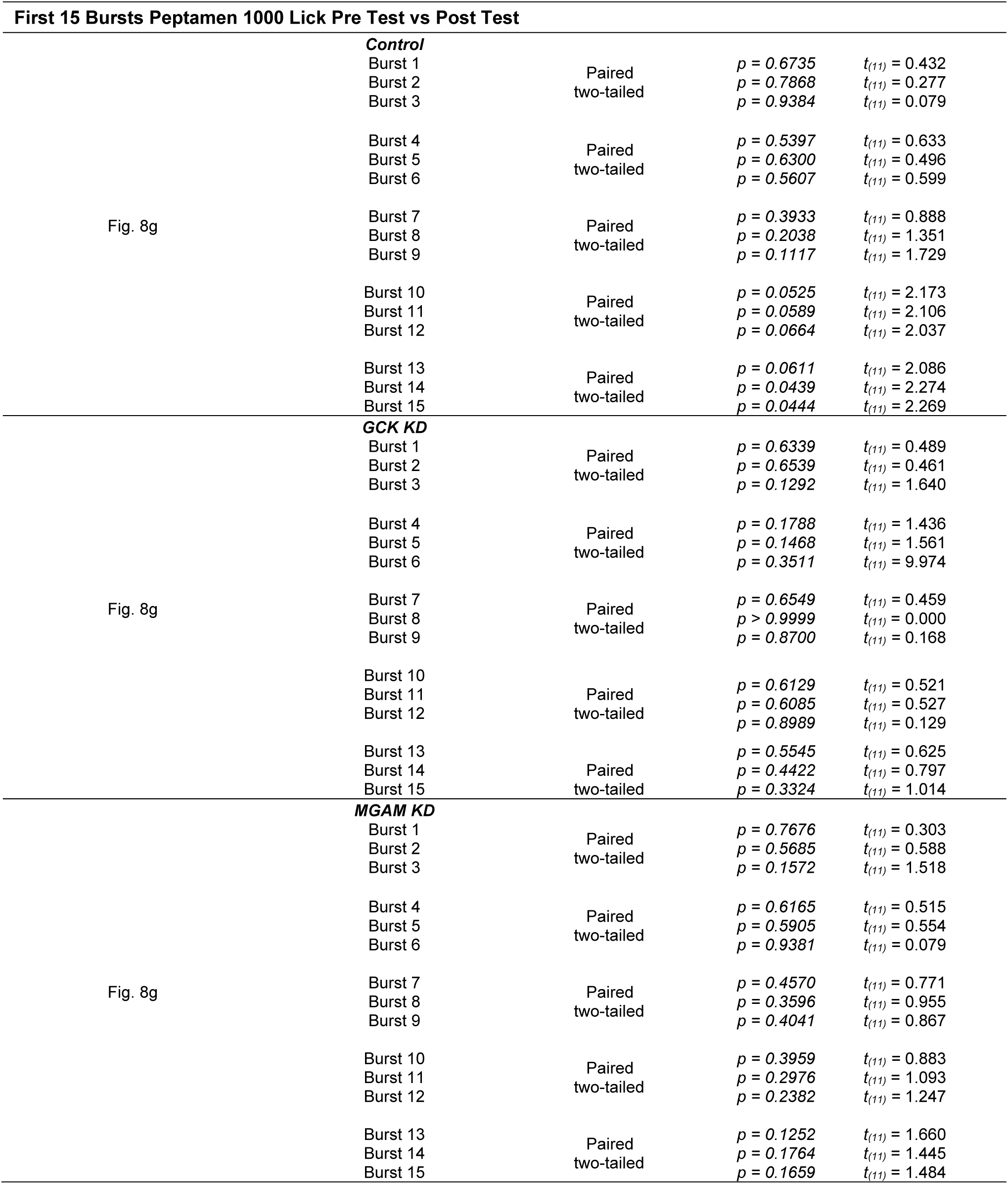
Student’s *t*-tests.

| Peptamen 1000 Lick Pre Test vs Post Test |  |  |  |  |
| --- | --- | --- | --- | --- |
| Fig. 8c | **Licks Used** |  |  |  |
| | Control | Paired<br>two-tailed | $p = 0.1105$ | $t_{(11)} = 1.735$ |
| | GCK KD | | $p = 0.1041$ | $t_{(11)} = 1.772$ |
| | MGAM KD | | $p = 0.0465$ | $t_{(11)} = 2.242$ |
| Fig. 8d | **Meal Duration** |  |  |  |
| | Control | Paired<br>two-tailed | $p = 0.0013$ | $t_{(11)} = 4.290$ |
| | GCK KD | | $p < 0.0001$ | $t_{(11)} = 8.166$ |
| | MGAM KD | | $p < 0.0001$ | $t_{(11)} = 9.330$ |
| Fig. 8e | **Burst Size** |  |  |  |
| | Control | Paired<br>two-tailed | $p = 0.0213$ | $t_{(11)} = 1.684$ |
| | GCK KD | | $p = 0.0465$ | $t_{(11)} = 2.243$ |
| | MGAM KD | | $p = 0.0251$ | $t_{(11)} = 2.590$ |
| Fig. 8f | **Burst Number** |  |  |  |
| | Control | Paired<br>two-tailed | $p = 0.0697$ | $t_{(11)} = 2.009$ |
| | GCK KD | | $p = 0.0115$ | $t_{(11)} = 3.029$ |
| | MGAM KD | | $p = 0.0542$ | $t_{(11)} = 2.154$ |
| First 15 Bursts Peptamen 1000 Lick Pre Test vs Post Test |  |  |  |  |
| Fig. 8g | **Control** |  |  |  |
| | Burst 1 | | $p = 0.6735$ | $t_{(11)} = 0.432$ |
| | Burst 2 | Paired | $p = 0.7868$ | $t_{(11)} = 0.277$ |
| | Burst 3 | two-tailed | $p = 0.9384$ | $t_{(11)} = 0.079$ |
| | Burst 4 | | $p = 0.5397$ | $t_{(11)} = 0.633$ |
| | Burst 5 | Paired | $p = 0.6300$ | $t_{(11)} = 0.496$ |
| | Burst 6 | two-tailed | $p = 0.5607$ | $t_{(11)} = 0.599$ |
| | Burst 7 | | $p = 0.3933$ | $t_{(11)} = 0.888$ |
| | Burst 8 | Paired | $p = 0.2038$ | $t_{(11)} = 1.351$ |
| | Burst 9 | two-tailed | $p = 0.1117$ | $t_{(11)} = 1.729$ |
| | Burst 10 | | $p = 0.0525$ | $t_{(11)} = 2.173$ |
| | Burst 11 | Paired | $p = 0.0589$ | $t_{(11)} = 2.106$ |
| | Burst 12 | two-tailed | $p = 0.0664$ | $t_{(11)} = 2.037$ |
| | Burst 13 | | $p = 0.0611$ | $t_{(11)} = 2.086$ |
| | Burst 14 | Paired | $p = 0.0439$ | $t_{(11)} = 2.274$ |
| | Burst 15 | two-tailed | $p = 0.0444$ | $t_{(11)} = 2.269$ |
| Fig. 8g | **GCK KD** |  |  |  |
| | Burst 1 | | $p = 0.6339$ | $t_{(11)} = 0.489$ |
| | Burst 2 | Paired | $p = 0.6539$ | $t_{(11)} = 0.461$ |
| | Burst 3 | two-tailed | $p = 0.1292$ | $t_{(11)} = 1.640$ |
| | Burst 4 | | $p = 0.1788$ | $t_{(11)} = 1.436$ |
| | Burst 5 | Paired | $p = 0.1468$ | $t_{(11)} = 1.561$ |
| | Burst 6 | two-tailed | $p = 0.3511$ | $t_{(11)} = 9.974$ |
| | Burst 7 | | $p = 0.6549$ | $t_{(11)} = 0.459$ |
| | Burst 8 | Paired | $p > 0.9999$ | $t_{(11)} = 0.000$ |
| | Burst 9 | two-tailed | $p = 0.8700$ | $t_{(11)} = 0.168$ |
| | Burst 10 | | $p = 0.6129$ | $t_{(11)} = 0.521$ |
| | Burst 11 | Paired | $p = 0.6085$ | $t_{(11)} = 0.527$ |
| | Burst 12 | two-tailed | $p = 0.8989$ | $t_{(11)} = 0.129$ |
| | Burst 13 | | $p = 0.5545$ | $t_{(11)} = 0.625$ |
| | Burst 14 | Paired | $p = 0.4422$ | $t_{(11)} = 0.797$ |
| | Burst 15 | two-tailed | $p = 0.3324$ | $t_{(11)} = 1.014$ |
| Fig. 8g | **MGAM KD** |  |  |  |
| | Burst 1 | | $p = 0.7676$ | $t_{(11)} = 0.303$ |
| | Burst 2 | Paired | $p = 0.5685$ | $t_{(11)} = 0.588$ |
| | Burst 3 | two-tailed | $p = 0.1572$ | $t_{(11)} = 1.518$ |
| | Burst 4 | | $p = 0.6165$ | $t_{(11)} = 0.515$ |
| | Burst 5 | Paired | $p = 0.5905$ | $t_{(11)} = 0.554$ |
| | Burst 6 | two-tailed | $p = 0.9381$ | $t_{(11)} = 0.079$ |
| | Burst 7 | | $p = 0.4570$ | $t_{(11)} = 0.771$ |
| | Burst 8 | Paired | $p = 0.3596$ | $t_{(11)} = 0.955$ |
| | Burst 9 | two-tailed | $p = 0.4041$ | $t_{(11)} = 0.867$ |
| | Burst 10 | | $p = 0.3959$ | $t_{(11)} = 0.883$ |
| | Burst 11 | Paired | $p = 0.2976$ | $t_{(11)} = 1.093$ |
| | Burst 12 | two-tailed | $p = 0.2382$ | $t_{(11)} = 1.247$ |
| | Burst 13 | | $p = 0.1252$ | $t_{(11)} = 1.660$ |
| | Burst 14 | Paired | $p = 0.1764$ | $t_{(11)} = 1.445$ |
| | Burst 15 | two-tailed | $p = 0.1659$ | $t_{(11)} = 1.484$ |

**Supplementary Table 8B:**
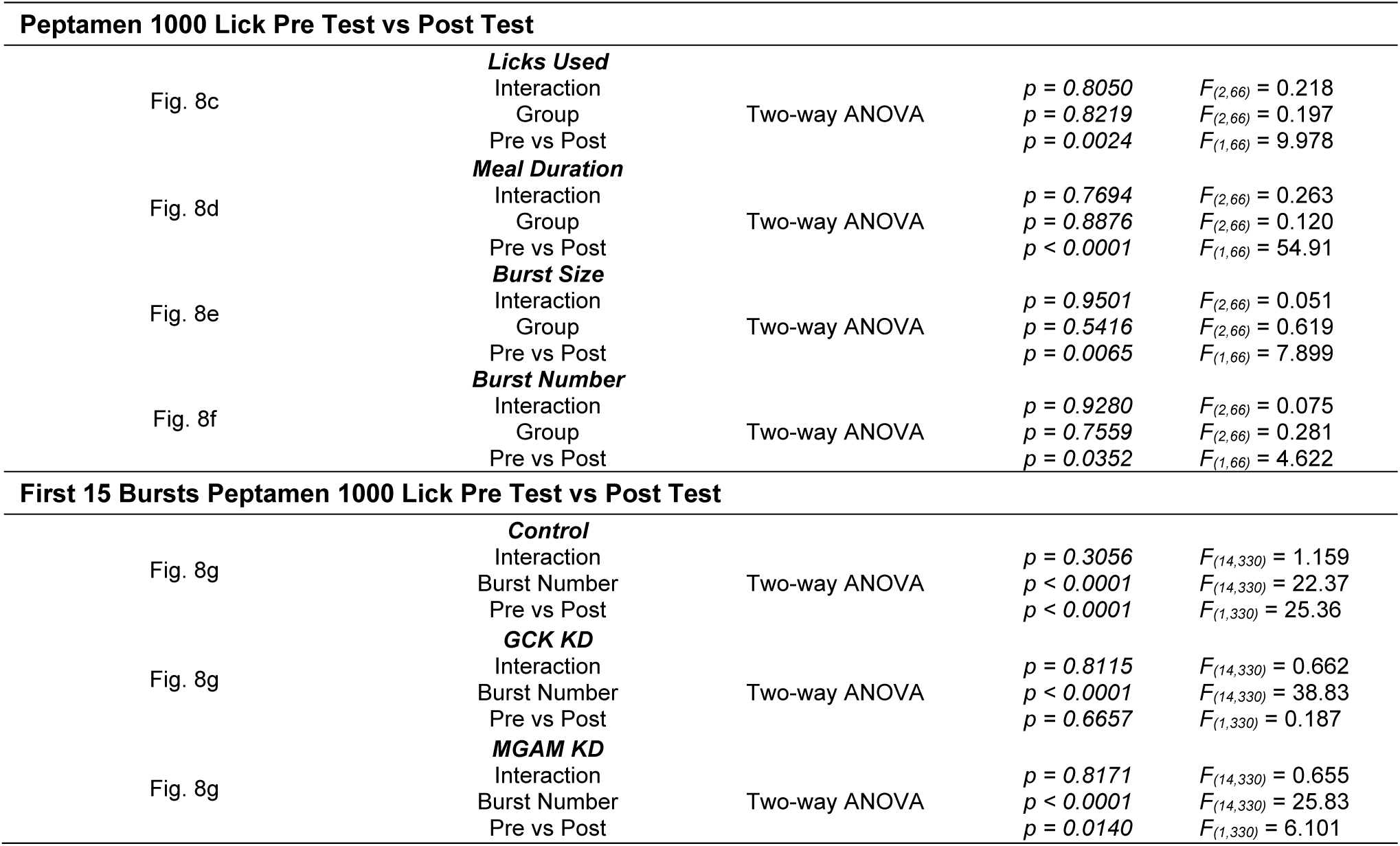
ANOVA tests.

**Supplementary Table 9A:**
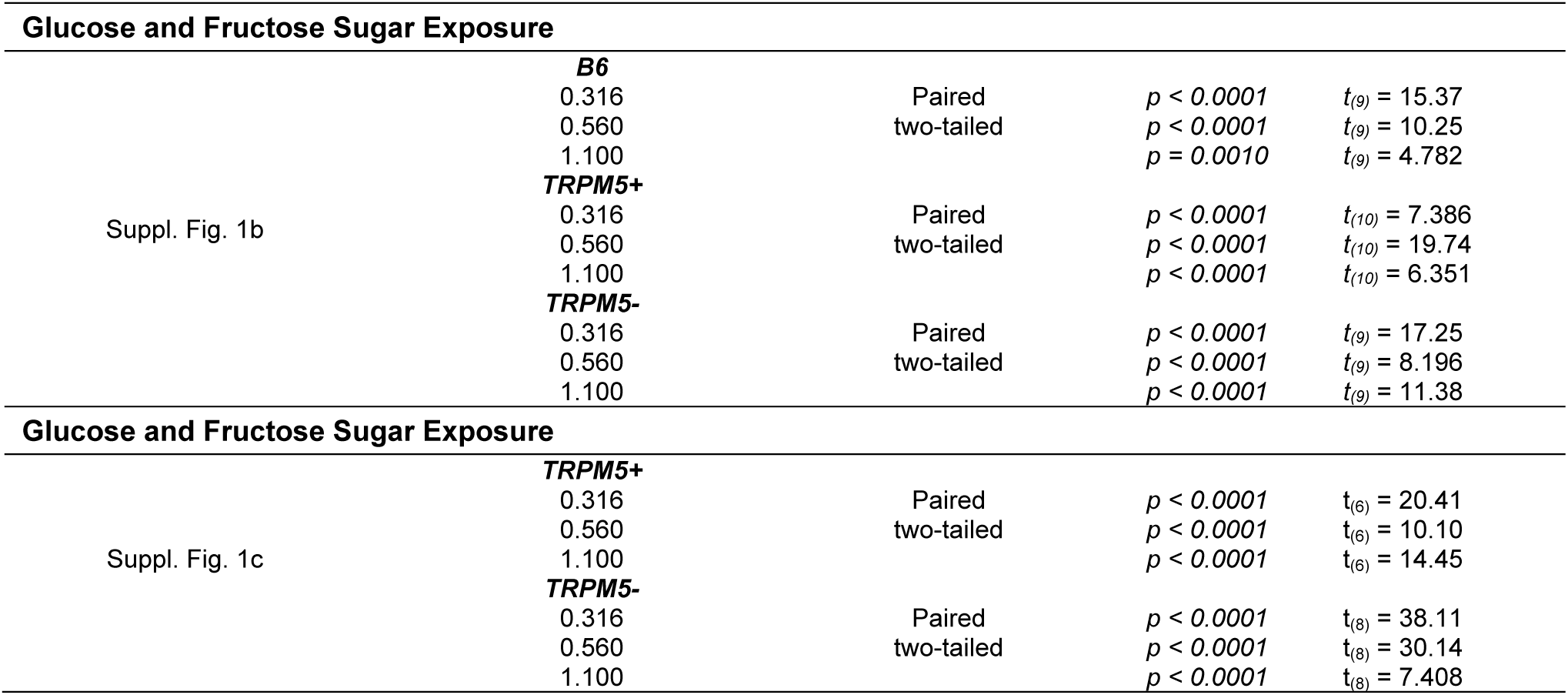
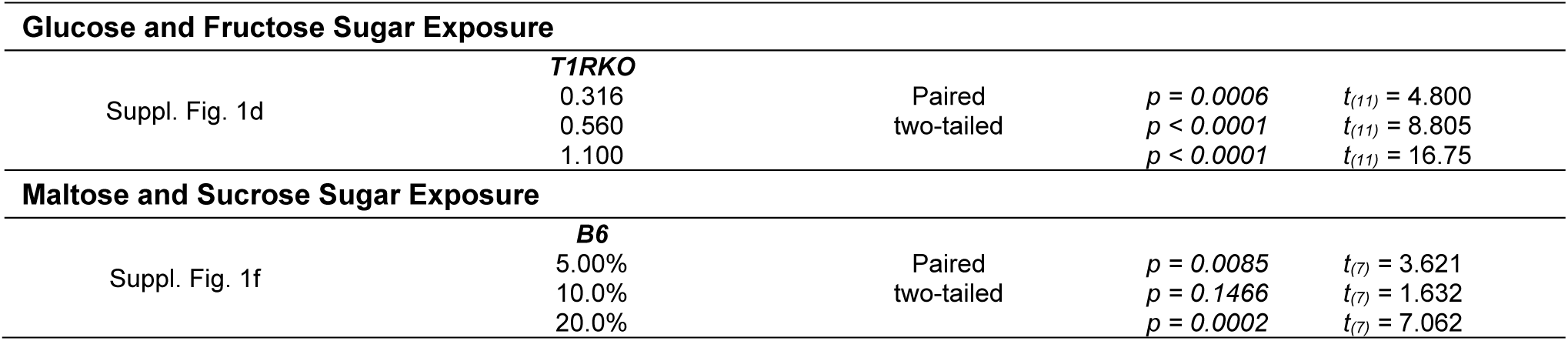
Student’s *t*-tests.

**Supplementary Table 9B:**
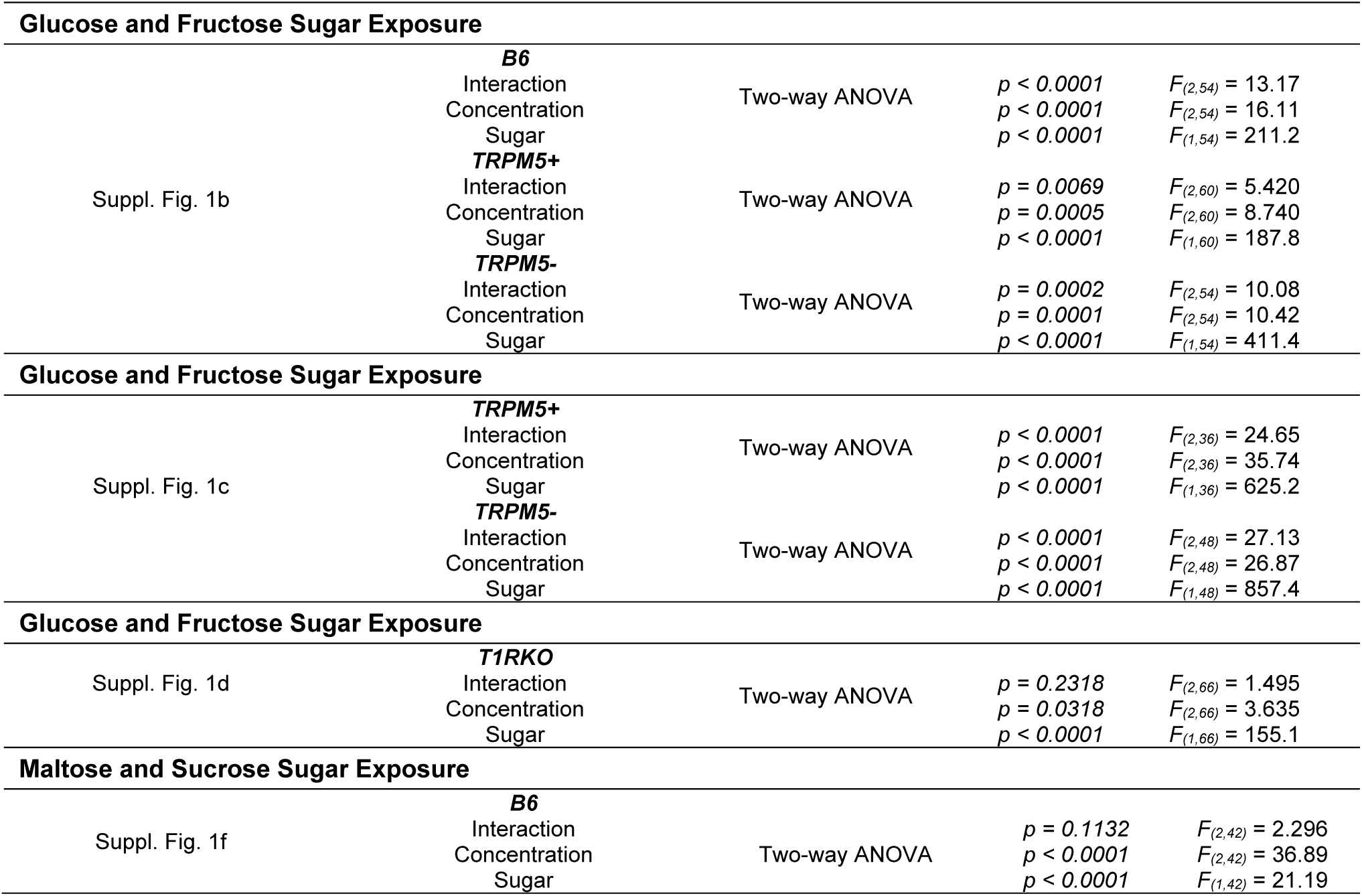
ANOVA tests.

**Supplementary Table 10A:** Student’s *t*-tests.

| Taste Genes mRNA Expression Levels |  |  |  |  |
| --- | --- | --- | --- | --- |
| Suppl. Fig. 2 | ***Trpm5* mRNA expression** |  |  |  |
| | SN B6 vs SE B6 | Unpaired<br>two-tailed | $p = 0.8788$ | $t_{(14)} = 1.553$ |
| | SN TRPM5+ vs SE TRPM5+ | | $p = 0.8327$ | $t_{(9)} = 0.217$ |
| | SN TRPM5- vs SE TRPM5- | | $p = 0.0532$ | $t_{(11)} = 2.166$ |
| | B6 and TRPM5+ vs TRPM5- | | $p < 0.0001$ | $t_{(38)} = 5.057$ |
|  | ***Tas1r3* mRNA expression** |  |  |  |
| | SN B6 vs SE B6 | Unpaired<br>two-tailed | $p = 0.6425$ | $t_{(14)} = 0.474$ |
| | SN TRPM5+ vs SE TRPM5+ | | $p = 0.4746$ | $t_{(9)} = 0.746$ |
| | SN TRPM5- vs SE TRPM5- | | $p = 0.3356$ | $t_{(11)} = 1.007$ |
| | B6 and TRPM5+ vs TRPM5- | | $p = 0.0445$ | $t_{(38)} = 2.078$ |
|  | ***Gck* mRNA expression** |  |  |  |
| | SN B6 vs SE B6 | Unpaired<br>two-tailed | $p = 0.6146$ | $t_{(14)} = 0.515$ |
| | SN TRPM5+ vs SE TRPM5+ | | $p = 0.6267$ | $t_{(8)} = 0.506$ |
| | SN TRPM5- vs SE TRPM5- | | $p = 0.2681$ | $t_{(11)} = 1.166$ |
| | B6 and TRPM5+ vs TRPM5- | | $p = 0.0132$ | $t_{(37)} = 2.604$ |
|  | ***Mgam* mRNA expression** |  |  |  |
| | SN B6 vs SE B6 | Unpaired<br>two-tailed | $p = 0.3318$ | $t_{(10)} = 1.020$ |
| | SN TRPM5+ vs SE TRPM5+ | | $p = 0.8312$ | $t_{(7)} = 0.221$ |
| | SN TRPM5- vs SE TRPM5- | | $p = 0.9405$ | $t_{(7)} = 0.077$ |
| | B6 vs TRPM5+ and TRPM5- | | $p = 0.0002$ | $t_{(28)} = 4.271$ |

**Supplementary Table 10B:** ANOVA tests.

| Taste Genes mRNA Expression Levels |  |  |  |  |
| --- | --- | --- | --- | --- |
| Suppl. Fig. 2 | *Trpm5* mRNA expression | One-way ANOVA | $p = 0.0004$ | $F_{(5,34)} = 6.121$ |
| | *Tas1r3* mRNA expression | One-way ANOVA | $p = 0.0718$ | $F_{(5,34)} = 2.248$ |
| | *Gck* mRNA expression | One-way ANOVA | $p = 0.0913$ | $F_{(5,33)} = 2.092$ |
| | *Mgam* mRNA expression | One-way ANOVA | $p = 0.0183$ | $F_{(5,24)} = 3.400$ |

**Supplementary Table 11A:** Student’s *t*-tests.

| Maltose vs Sucrose Brief Access Taste Test |  |  |  |  |
| --- | --- | --- | --- | --- |
| Suppl. Fig. 3a | Control |  |  |  |
| | 0.316 | Paired two-tailed | $p = 0.0564$ | $t_{(4)} = 2.660$ |
| | 0.560 | | $p = 0.3824$ | $t_{(4)} = 0.981$ |
| | 1.100 | | $p = 0.0724$ | $t_{(4)} = 2.424$ |
|  | MGAM KD |  |  |  |
| | 0.316 | Paired two-tailed | $p = 0.8806$ | $t_{(4)} = 0.160$ |
| | 0.560 | | $p = 0.4365$ | $t_{(4)} = 0.864$ |
| | 1.100 | | $p = 0.2360$ | $t_{(4)} = 1.393$ |
| Average Maltose vs Sucrose Brief Access Taste Test |  |  |  |  |
| Suppl. Fig. 3b | Control Maltose vs Sucrose | Paired two-tailed | $p = 0.2144$ | $t_{(4)} = 1.474$ |
| | MGAM KD Maltose vs Sucrose | Paired two-tailed | $p = 0.8823$ | $t_{(4)} = 0.158$ |
| | Control Maltose vs GCK KD Maltose | Unpaired two-tailed | $p = 0.6629$ | $t_{(8)} = 0.453$ |
| | Control Sucrose vs GCK KD Sucrose | Unpaired two-tailed | $p = 0.9958$ | $t_{(8)} = 0.005$ |
| Glucose vs Fructose Brief Access Taste Test |  |  |  |  |
| Suppl. Fig. 3c | Control |  |  |  |
| | 0.316 | Paired two-tailed | $p = 0.0149$ | $t_{(5)} = 3.640$ |
| | 0.560 | | $p = 0.0030$ | $t_{(5)} = 5.392$ |
| | 1.100 | | $p = 0.0005$ | $t_{(5)} = 7.998$ |
|  | MGAM KD |  |  |  |
| | 0.316 | Paired two-tailed | $p = 0.0338$ | $t_{(4)} = 3.172$ |
| | 0.560 | | $p = 0.2428$ | $t_{(4)} = 1.369$ |
| | 1.100 | | $p = 0.0004$ | $t_{(4)} = 11.00$ |
| Average Glucose vs Fructose Brief Access Taste Test |  |  |  |  |
| Suppl. Fig. 3d | Control Glucose vs Fructose | Paired two-tailed | $p < 0.0001$ | $t_{(5)} = 14.60$ |
| | MGAM KD Glucose vs Fructose | Paired two-tailed | $p = 0.0102$ | $t_{(4)} = 4.575$ |
| | Control Glucose vs MGAM KD Glucose | Unpaired two-tailed | $p = 0.8485$ | $t_{(9)} = 0.197$ |

**Supplementary Table Sugar 11B:**
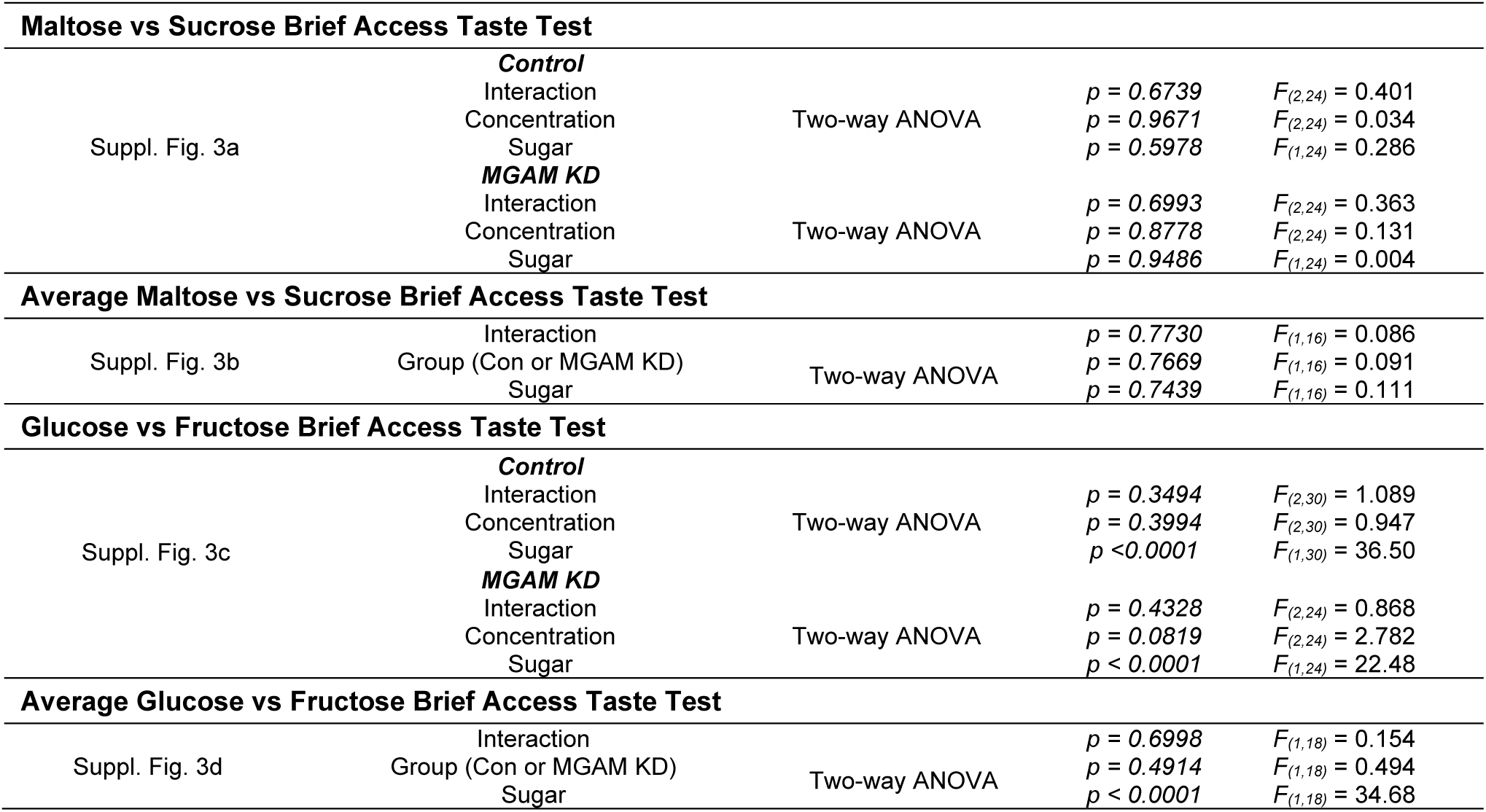
ANOVA tests.

